# Gamete-specific expression of TALE class HD genes activates the diploid sporophyte program in *Marchantia polymorpha*

**DOI:** 10.1101/2020.04.06.027821

**Authors:** Tom Dierschke, Eduardo Flores-Sandoval, Madlen I. Rast-Somssich, Felix Althoff, Sabine Zachgo, John L. Bowman

## Abstract

Eukaryotic life cycles alternate between haploid and diploid phases and in phylogenetically diverse unicellular eukaryotes, expression of paralogous homeodomain genes in the two gametes directs the haploid-to-diploid transition. In the unicellular Chlorophyte alga *Chlamydomonas* KNOX and BELL TALE-homeodomain genes mediate the transition. Here we demonstrate that in the liverwort *Marchantia polymorpha* paternal (sperm) expression three of the five phylogenetically diverse BELL genes, Mp*BELL234*, and maternal (egg) expression of Mp*KNOX1* mediate the haploid-to-diploid transition. Loss-of-function alleles of either result in zygotic or early embryonic arrest. In land plants both the haploid gametophyte and diploid sporophyte are complex multicellular organisms. Expression of Mp*KNOX1* and two other paralogs, Mp*BELL1* and Mp*KNOX2*, during sporophyte development is consistent with a later role in patterning the sporophyte. These results indicate that the ancestral mechanism to activate diploid gene expression was retained in early diverging land plants and subsequently co-opted during evolution of the diploid sporophyte body.

## Introduction

The life cycles of eukaryotes alternate between haploid and diploid phases, initiated by meiosis and gamete fusion, respectively. Expression of paralogous homeodomain genes in the two gametes directs the haploid-to-diploid transition in gene expression in phylogenetically diverse eukaryotes, including the ascomycete fungus *Saccharomyces cerevisiae* (Goutte and Johnson, 1988, Herskowitz, 1989), the basidiomycete fungi *Coprinopsis cinerea* and *Ustilago maydis* (Gillissen et al., 1992, Hull et al., 2005, Kues et al., 1992, Spit et al., 1998, Urban et al., 1996), the Amoebozoa *Dictyostelium discoideum* (Hedgethorne et al., 2017), the brown alga *Ectocarpus* (Arun et al., 2019), the red alga *Pyropia yezoensis* (Mikami et al., 2019), and the unicellular Chlorophyte alga *Chlamydomonas reinhardtii* (Ferris and Goodenough, 1987, Lee et al., 2008, Nishimura et al., 2012, Zhao et al., 2001). This broad phylogenetic distribution suggest this was an ancestral function of homeodomain genes [reviewed in (Bowman et al., 2016b)]. In the Viridiplantae the paralogous genes are two subclasses, KNOX and BELL, of TALE-class homeodomain (HD) genes. In *Chlamydomonas*, the minus (-) gamete expresses a KNOX protein (GSM1) and the plus (+) gamete expresses a BELL protein (GSP1), and upon gamete fusion the two proteins heterodimerize and translocate to the nucleus activating zygotic gene expression (Lee et al., 2008). GSM1 and GSP1 are necessary for diploid gene expression, and when ectopically expressed together in vegetative haploid cells are sufficient to induce the diploid genetic program (Ferris and Goodenough, 1987, Lee et al., 2008, Nishimura et al., 2012, Zhao et al., 2001). Biologically, the expression of a unique paralog in each type of gamete, coupled with the requirement for heterodimerization for functionality, is a mechanism to preload gametes such that immediately following gamete fusion/fertilization a distinct diploid genetic program is initiated.

Heterodimerization of TALE-HD paralogs is mediated by subclass-specific protein domains N-terminal of the homeodomain that are in some cases conserved in phylogenetically disparate eukaryotes (Bellaoui et al., 2001, Burglin, 1997). For example, Viridiplantae KNOX proteins and metazoan MEIS proteins share a homologous heterodimerization domain thus defining a TALE-HD class, named MEINOX, present in the ancestral eukaryote (Burglin, 1997). In contrast, heterodimerization domains of MEINOX partners are not as well conserved (Joo et al., 2018). Some MEINOX partners in animals have a characteristic PBC-C heterodimerization domain (Burglin, 1997), and PBC-related domains are found in some algal BELL-related TALE-HD proteins, suggesting its presence in the ancestral eukaryote (Joo et al., 2018). In contrast, land plant BELL proteins possess conserved SKY and BELL domains, collectively termed POX (pre-homeobox), that mediate heterodimerization with KNOX partners (Bellaoui et al., 2001, Hackbusch et al., 2005, Smith et al., 2002). POX domains are either highly diverged from, or unrelated to, PBC domains. Finally, in other algal BELL-related TALE-HD proteins neither heterodimerization domain is evident (Joo et al., 2018). Regardless, when tested for dimerization potential, Archaeplastida BELL-related TALE-HD proteins interact only with MEINOX TALE-HD partners and not with other BELL-related TALE-HD proteins (Joo et al., 2018).

Land plants are characterized by an alternation of generations whereby complex multicellular bodies develop in both haploid (gametophyte) and diploid (sporophyte) phases of the life cycle (Hofmeister, 1862) and elements of the KNOX/BELL system have been implicated in regulating the land plants haploid to diploid transition. In the moss *Physcomitrella patens*, one subclass of KNOX genes, KNOX1, is required for proper proliferation and differentiation in the diploid body (Sakakibara et al., 2008, Singer and Ashton, 2007), while another subclass, KNOX2, acts to suppress the haploid genetic program during diploid development (Sakakibara et al., 2013). In addition to being expressed during sporophyte development, both classes of KNOX genes are expressed in the egg cell during gamete formation (Sakakibara et al., 2013, Sakakibara et al., 2008). The expression of KNOX genes in egg cells suggests that this gamete may correspond to the (-) gamete in *Chlamydomonas*, which also expresses a KNOX protein, implying that the male gamete in land plants might express BELL proteins, as does the (+) gamete in *Chlamydomonas* (Sakakibara et al., 2013, Sakakibara et al., 2008). Consistent with this hypothesis, ectopic expression of a *Physcomitrella* BELL paralog, Pp*BELL1*, has been noted to induce the diploid genetic program in specific cell types of the gametophyte, but Pp*BELL1* was paradoxically reported to be expressed in the egg cell rather than the male gamete as might have been expected (Horst et al., 2016). However, expression of Pp*BELL1* has also been reported to be induced by activation of glutamate channels in the sperm, with Pp*BELL1* subsequently functionally active in the sporophyte (Ortiz-Ramírez et al., 2017). Finally, in *Physcomitrella* the KNOX/BELL diploid genetic program is actively suppressed during vegetative haploid development via polycomb-mediated repression (Mosquna et al., 2009, Okano et al., 2009). These observations prompted our investigation of the homologous genetic programs in the liverwort *Marchantia polymorpha*.

## Results

### *M. polymorpha* possesses phylogenetically diverse TALE homeodomain proteins

The *M. polymorpha* genome encodes nine TALE-HD-related family members: four KNOX genes and five BELL genes [(Bowman et al., 2017); **Figure 1**], all of which are expressed in the haploid sexual organs or diploid sporophyte, with minimal or no expression detected by rtPCR in the haploid vegetative thallus (**Figure 1—figure supplement 1**). Of the four KNOX genes, three are KNOX1 subclass, with only one encoding a HD, and one KNOX2 subclass (**Figure 1—figure supplement 1**). The two subclasses arose via gene duplication in an ancestral charophycean alga (Sakakibara, 2016, Joo et al., 2018, Frangedakis et al., 2017). Expression of Mp*KNOX1* (Mp5g01600/Mapoly0175s0020), which encodes a HD, is predominantly detected in archegoniophores and young sporophytes. In contrast, the KNOX1 genes lacking an HD (Mp*KNOX1A*, Mp4g12450/Mapoly0174s0007; Mp*KNOX1B*, Mp2g11140/Mapoly0023s0081) are predominantly expressed in antheridiophores. Mp*KNOX2* (Mp7g05320/Mapoly0194s0001) is expressed primarily during sporophyte development.

**Figure 1.**
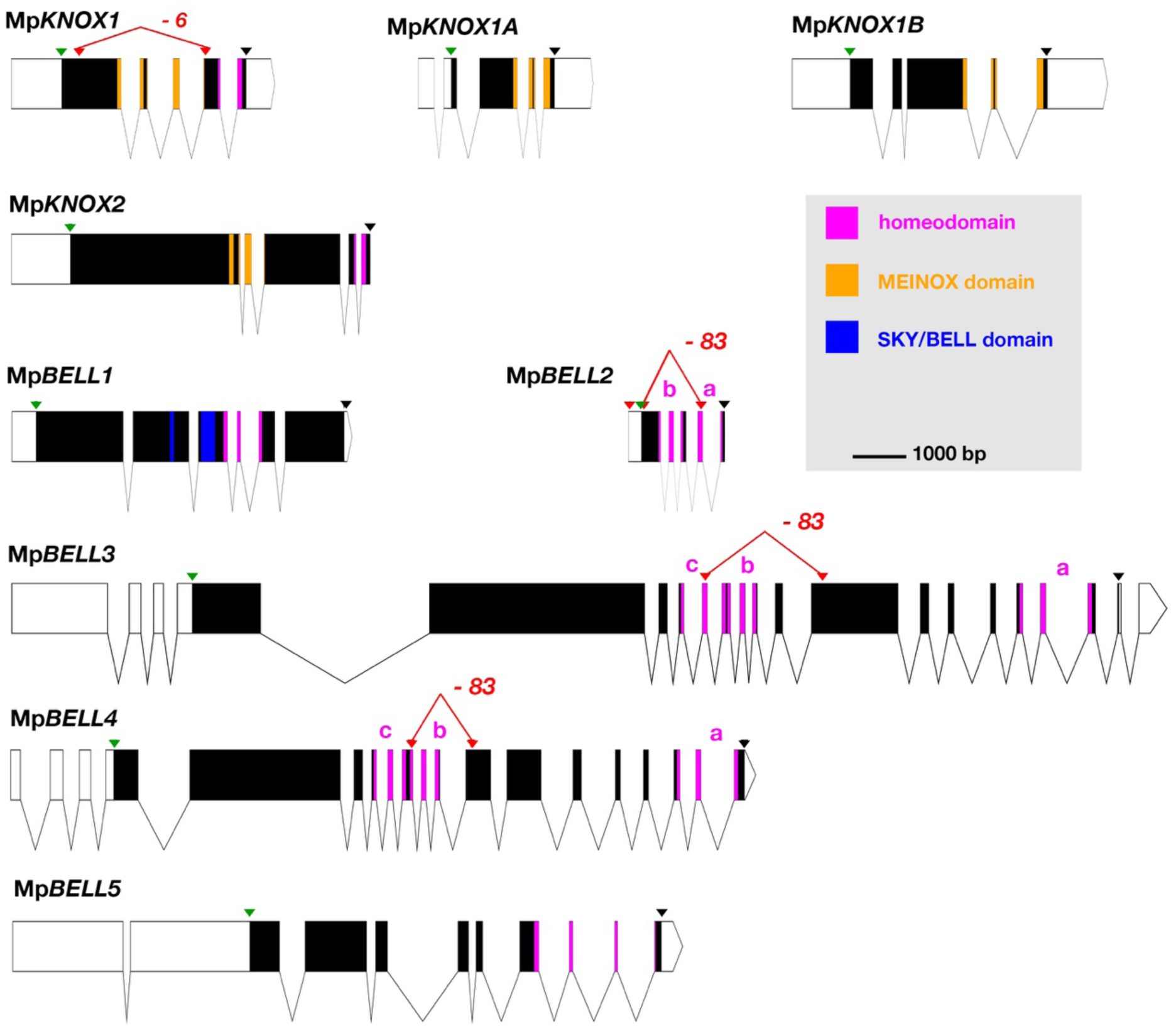
Schematic representations of the nine TALE-HD loci of *M. polymorpha*. White, 5’ and 3’ UTRs; black, coding exons; thin lines, introns; green triangles, start codon; black triangles, stop codon; red triangles, guide RNA targeted positions. The mutant alleles generated via CRISPR-Cas9 are described in more detail in Table 1. All protein annotation models are based on the *Marchantia* genome assembly of v5.1 except for Mp*KNOX2*. The Mp*KNOX2* model is based on sequences derived from RT-PCR. In genes with multiple homeodomains, they are denoted a, b and c. Gene models were assembled using wormweb (http://wormweb.org/exonintron).

**Table 1.**
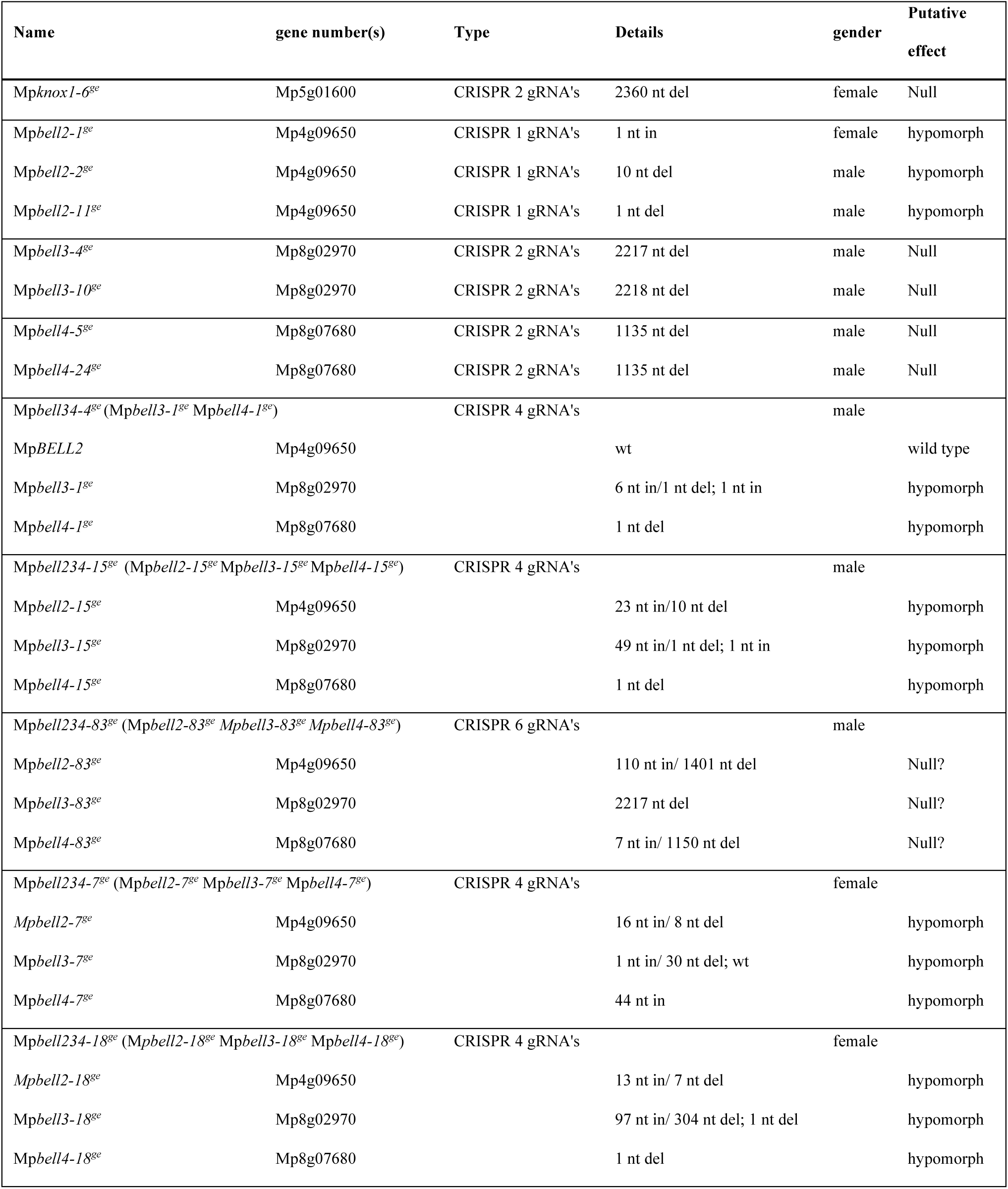
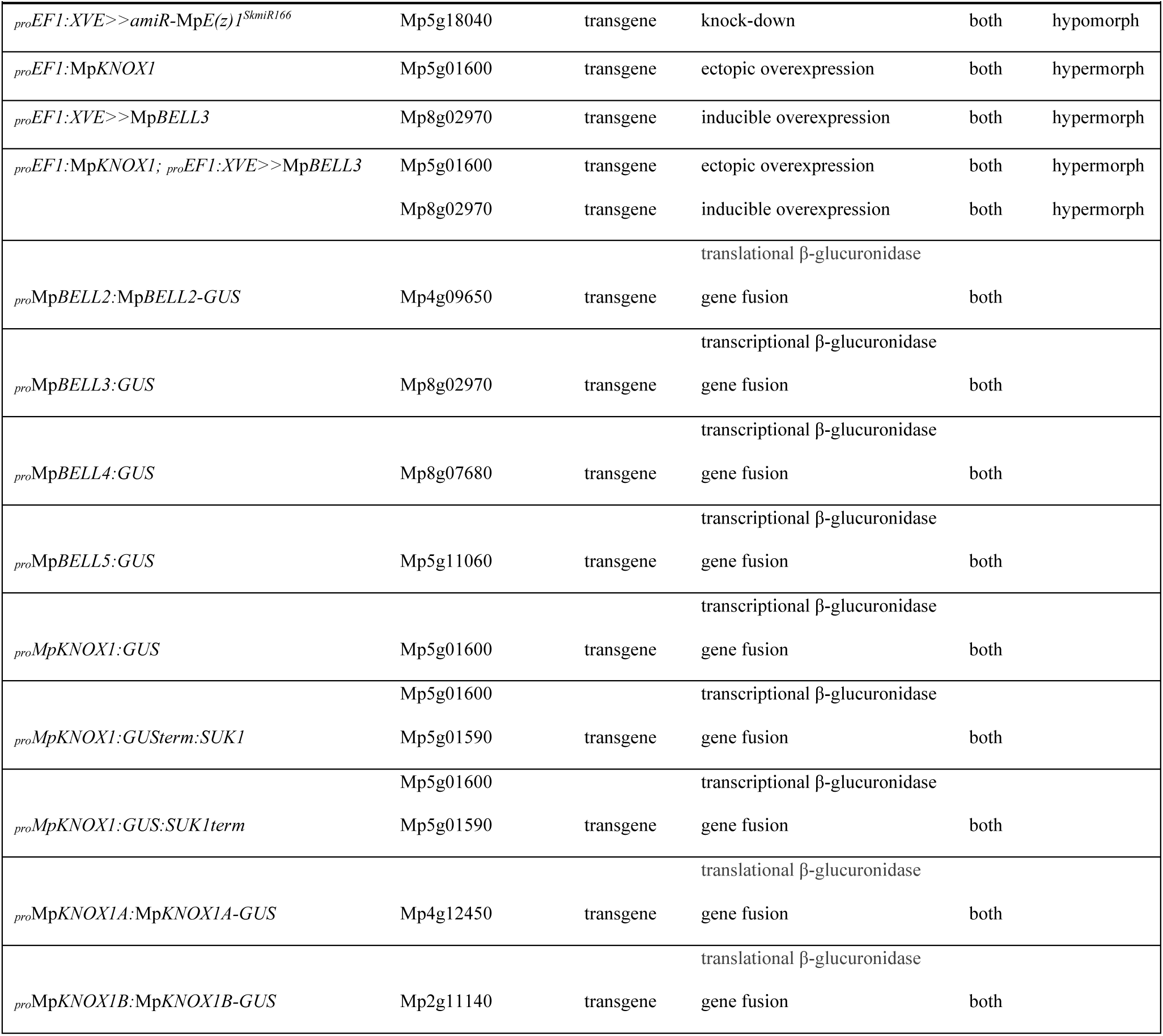
Alleles used in this study. Nomenclature is as outlined previously (Bowman et al., 2016a, Montgomery et al., 2020).

In contrast to the *M. polymorpha* KNOX genes, only one BELL gene (Mp*BELL1*, Mp8g18310/Mapoly0213s0014) is phylogenetically related to previously described land plant BELL genes, whereas the other four are more closely related to algal TALE-HD genes (**Figure 2**). Mp*BELL1* harbors a conserved canonical land plant BELL homeodomain sequence and a discernible, albeit divergent, BELL domain characteristic of other land plant BELL genes. Mp*BELL1* expression is detected predominantly during sporophyte development (**Figure 1— figure supplement 1**). Three of the algal-like BELL genes, [Mp*BELL2* (Mp4g09650/Mapoly0132s0008), Mp*BELL3* (Mp8g02970/Mapoly0012s0090) and Mp*BELL4* (Mp8g07680/Mapoly0013s0027)], each encode two or three homeodomains (**Figure 1**). The carboxyl HD is most conserved and those upstream becoming progressively less conserved consistent with their origins being via successive intragenic duplications (**Figure 2—figure supplement 1**). Notably, Mp*BELL*3 and Mp*BELL4* both encode large proteins and are expressed primarily in antheridiophores, with Mp*BELL2* only encoding the two HDs. These three genes are related via gene duplications within liverworts, but their phylogenetic affinity to other algal BELL classes is enigmatic (**Figure 2**). The fourth algal-like sequence, Mp*BELL5* (Mp5g11060/Mapoly0093s0028), is phylogenetically distinct, and resides in the previously defined GLX-basal clade (Joo et al., 2018). Unlike some algal BELL-related proteins (Joo et al., 2018), the presence of a PBC domain is not easily discernible in any of the predicted *M. polymorpha* BELL protein sequences. Thus, a diversity of BELL-related paralogs arose in an algal ancestor and has persisted in *M. polymorpha*. The presence of an algal-related *Metzgeria* BELL sequence implies that BELL diversity may exist throughout liverworts.

**Figure 2.**
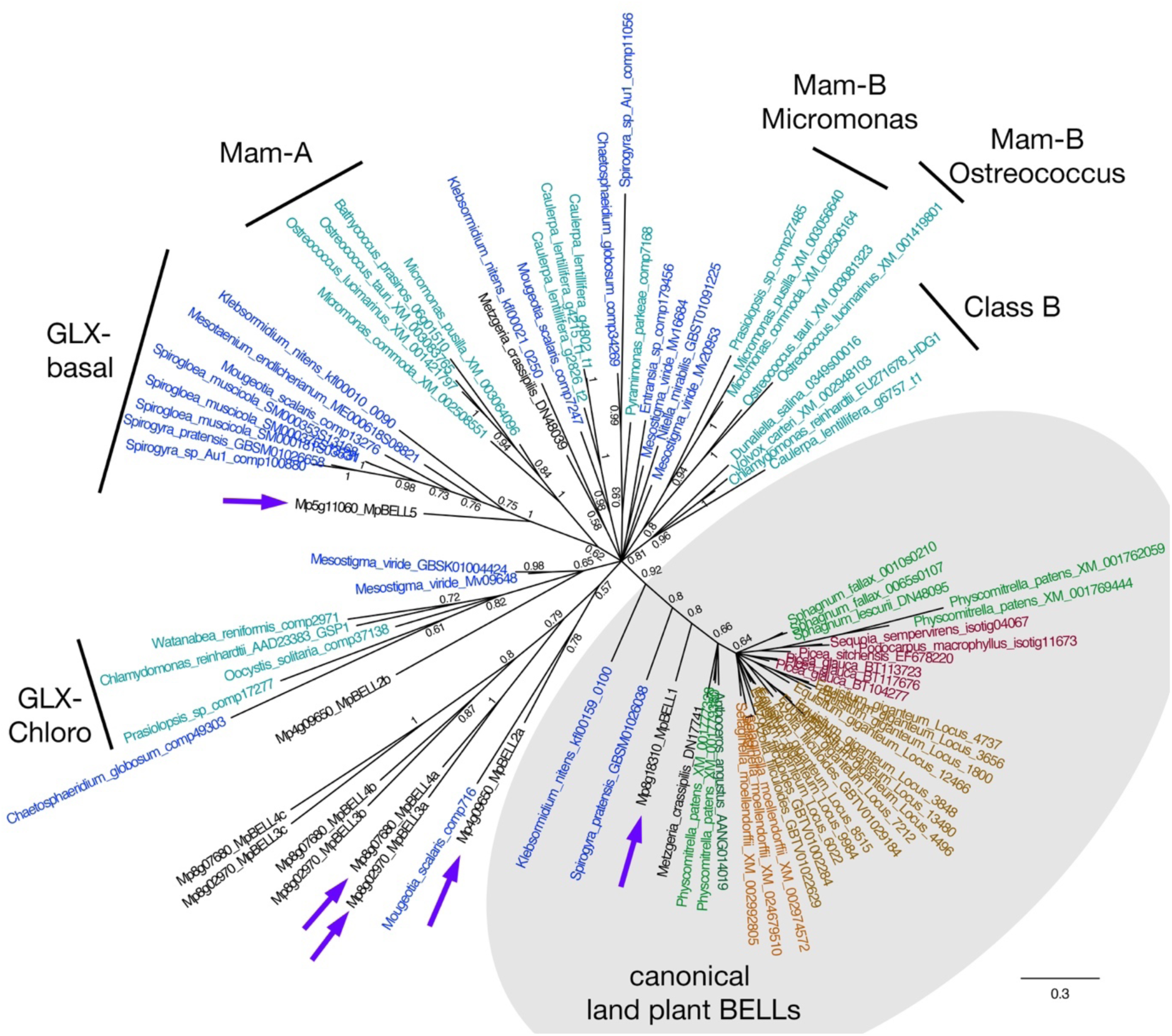
Unrooted Bayesian phylogram of Viridiplantae BELL-TALE class homeodomain genes. Tree was constructed using a nucleotide alignment of the homeodomain; a phylogenetic tree constructed using amino acid characters was congruent, albeit with less resolution. The five *M. polymorpha* BELL genes are highlighted (purple arrows). A clade of genes representing land plant BELL genes previously identified in land plants (gray shading) is recovered with high support. This clade includes Mp*BELL1* and charophycean algal sequences dating as far back as prior to the divergence of *Klebsormidium*. The canonical land plant BELL genes also share significant sequence similarity outside the homeodomain, including the SKY and BELL domains. The remaining BELL genes, including all Chlorophyte sequences, sequences from charophycean algae, and the other four *M. polymorpha* sequences reside in an unresolved polytomy. Within the unresolved polytomy, several subclades (labelled) previously identified are evident (Joo et al., 2018). The multiple homeodomains of Mp*BELL2*, Mp*BELL3* and Mp*BELL4* are designated a, b and c (Figure 1). Chlorophyte, light blue; charophyte, dark blue; liverwort, black; moss, green; hornwort, dark green; lycophyte, brown, fern, orange; seed plant, red. Numbers at branches indicate posterior probability values >50%; values within the polytomy in the canonical land plant clade were omitted for clarity.

### Vegetative gametophytic knock-down of Mp*E(z)* de-represses Mp*KNOX2* and Mp*BELL1* expression

In the moss *P. patens,* genome wide polycomb repressive complex 2 (PRC2)-mediated repression of gene expression is observed in the shift from gametophyte to sporophyte and at least a subset of KNOX and BELL TALE-HD genes exhibit polycomb-mediated repression in the *P. patens* vegetative gametophyte (Mosquna et al., 2009, Okano et al., 2009, Pereman et al., 2016). Thus, we first examined whether any *Marchantia* TALE-HD genes are repressed in the vegetative gametophyte in a PRC2-dependent manner.

PRC2, found in both animals and plants (Goodrich et al., 1997, Jones and Gelbart, 1993), acts as an epigenetic regulator of genetic programs, and thus provides a ‘cellular memory’ (Cao et al., 2002, Czermin et al., 2002, Saurin et al., 2001). Each PRC2 includes a member of the SET domain family, which has methyltransferase activity that enzymatically methylates histones (Fischle et al., 2003), with trimethylation of lysine 27 at histone 3 (H3K27me3) being a genomic imprint of PRC2 that is inherited throughout a cell lineage (Cao et al., 2002, Czermin et al., 2002, Muller et al., 2002). The *M. polymorpha* genome harbors three *Enhancer of zeste* orthologs, Mp*E(z)1* (Mp5g18040/Mapoly0084s00510), Mp*E(z)2* (Mp5g17740/Mapoly0084s0024) and Mp*E(z)3* (Mp5g22900/Mapoly0010s0166), that encode the histone methyltransferase catalytic subunit of PRC2. While Mp*E(z)1* is expressed in both gametophyte and sporophyte generations, expression of Mp*E(z)2* and Mp*E(z)3* is largely limited to the sporophyte (Bowman et al., 2017). These SET-domain containing methyltransferases act together with three other proteins in the PRC2. For each of the other members of the PRC2 only a single ortholog is present in *Marchantia*, and these other components are expressed in both generations (Bowman et al., 2017).

To examine Mp*E(z)1* function in the gametophyte, we designed an artificial miR *amiR-*Mp*E(z)1^SkmiR166^* to target Mp*E(z)1* transcripts. Since constitutive expression of the amiR was lethal, a strategy to enable inducible expression of *amiR-*Mp*E(z)1^SkmiR166^* was devised via an estrogen inducible system (*_pro_EF1:XVE*>>*amiR-*Mp*E(z)1^SkmiR166^*) allowing recovery of phenotypically wild-type plants (Flores-Sandoval et al., 2016). The down-regulation of Mp*E(z)1* should reduce PRC2 activity and hence reduce overall H3K27me3 marks within the genome, ‘opening up’ the chromatin for transcription of formerly repressed genes. To obtain an overview of the H3K27me3 chromatin landscape in *M. polymorpha* vegetative gametophytes, we sampled wild-type vegetative gametophytic tissue of 2-week-old male and female gemmae, as well as un-induced and induced male and female gemmae of *_pro_EF1:XVE>>amiR-*Mp*E(z)1*^SkmiR166^ lines of the same age. Compared to the overall H3K27me3 landscape in wild-type gametophytic tissue, the number and amplitude of methylation peaks are consistently reduced after 48h of down-regulation of Mp*E(z)1*.

We specifically examined the H3K27me3 pattern relative to whether gene expression was de-repressed at the nine TALE-HD loci in the *_pro_EF1:XVE*>>*amiR-*Mp*E(z)^SkmiR166^* background. Analysis of *_pro_EF1:XVE*>>*amiR-*Mp*E(z)^SkmiR166^* plants grown in the presence of 17-*β*-estradiol revealed that two TALE genes, Mp*KNOX2* and Mp*BELL1* (the canonical land plant BELL gene), are de-repressed (**Figure 3A**). De-repression of Mp*BELL1* was detectable after two hours while that of Mp*KNOX2* was detected after 6 hours, with expression of both increasing over 24 hours. Notably, these two genes, Mp*KNOX2* and Mp*BELL1* are the only two TALE-HD genes expressed largely during the stage of sporophyte development and not in the gametophytic reproductive structures (**Figure 3B, Figure 1—figure supplement 1**). Both Mp*KNOX2* and Mp*BELL1* also exhibited altered H3K27me3 patterns, being heavily marked in gametophytic wild-type tissue but losing most marks following induction as de-repression was observed (**Figure 3C**). Since neither Mp*KNOX2* nor Mp*BELL*1 expression was detected in the gametophytic sexual organs, where the egg cells and sperm are located, (**Figure 3C, Figure 1—supplement figure 1**), we considered these genes unlikely to play a direct role in the gametophyte to sporophyte transition.

**Figure 3.**
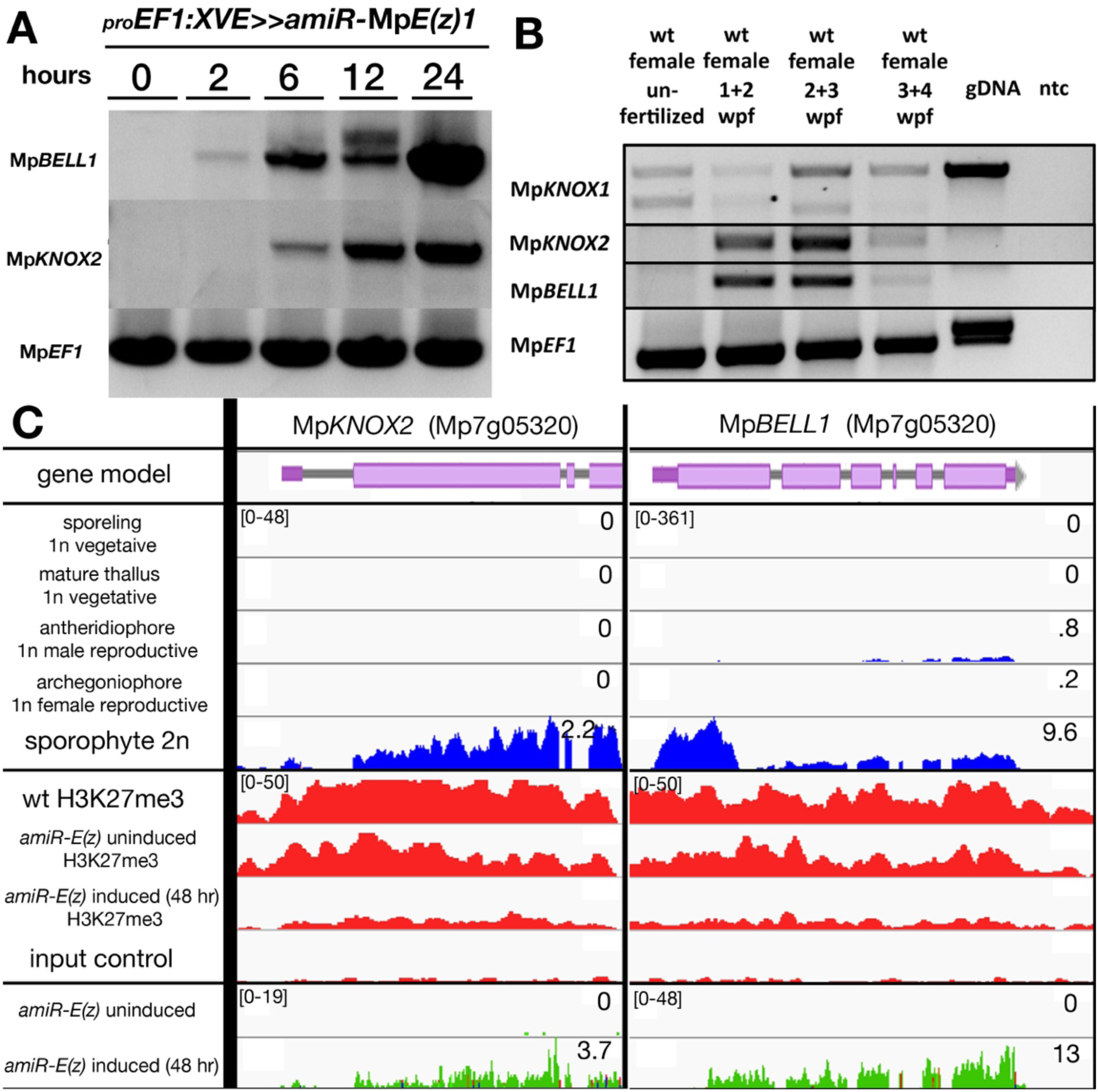
Reduction of Mp*E(z)1* activity results in de-repression of Mp*KNOX2* and Mp*BELL1*. (**A**) sqRT-PCR of gemmae grown on ½ B5 media containing 5μM 17-*β*-estradiol for said time points show expression of Mp*BELL1* and Mp*KNOX2* 2h and 6h post induction, respectively. Gene expression in vegetative gametophytes at different time points following induction on 5μM 17-β-estradiol plates. (**B**) Expression, as monitored by semi-quantitative RT-PCR, of Mp*KNOX1*, Mp*KNOX2*, and Mp*BELL1* in unfertilized archegonia, 1-2 week-old sporophytes, 2-3 week-old sporophytes and 3-4 week-old sporophytes. (**C**) Expression patterns and H3K27me3 marks at Mp*BELL1* and Mp*KNOX2*. Top panel is the predicted gene model. Panels in blue denote expression in different tissues of wild-type *M. polymorpha* based on RNA-seq experiments (Bowman et al., 2017). Panels in red show H3K27me3 patterns in vegetative thallus as determined by genome-wide ChIP-seq (see Materials and Methods). Panels in green are RNA-seq expression data in *_pro_EF1:XVE>>amiR-*Mp*E(z)^SkmiR166^* lines. Both Mp*KNOX2* and Mp*BELL1* genomic loci are marked with H3K27me3 in wild-type and uninduced lines. Both exhibit substantial reduction in H3K27me3 after 48h post induction and Mp*KNOX2* and Mp*BELL1* become de-repressed following induction. All scales (FPKM) are set to a standard within each outlined box; all H3K27me3 panels set to 50; for RNA-seq data the FPKM values (if less than 10, rounded to .1) are presented.

### Maternal Mp*KNOX1* is required for post-zygotic embryo development

During the gametophyte generation Mp*KNOX1* is expressed specifically in the egg cell, but is not detected at the stage prior when the venter canal cell is present (**Figure 4A-B**). During differentiation of the egg cell, Mp*KNOX1* appears to be expressed in a pulse early, with signal diminishing in older archegonia (**Figure 4B**). Following fertilization, Mp*KNOX1* is expressed throughout the developing sporophyte up until the time when future sporogenous cells become distinct (**Figure 4C-E**), with expression thereafter becoming undetectable during meiotic stages (**Figure 4F**). A female harboring a loss-of-function allele consisting of a 2.3 kb deletion spanning the Mp*KNOX1* locus (**Figure 1; Table 1**) was crossed with wild-type males. In contrast to control crosses using wild-type males and females where sporophyte production was observed (**Figure 4G-I**), mature sporophytes did not develop on crosses with the Mp*knox1*-6^ge^ females (**Figure 4J-L**). Closer examination of female Mp*knox1-6^ge^* mutants revealed a developmental arrest of the zygote following fertilization, with the zygote failing to undergo cytokinesis (**Figure 4K**). However, two gametophytic tissues, the calyptra and the pseudoperianth, whose growth is dependent upon fertilization (Hofmeister, 1862), commence develop normally (**Figure 4H-I, K-L**). Mp*knox1-6^ge^* mutants, as in wild-type, the calyptra undergoes periclinal divisions indicating successful fertilization and implying an intergenerational signal inducing its development. Likewise, the pseudoperianth, a ring of tissue surrounding the archegonium that only develops post-fertilization, initially develops normally in Mp*knox1-6^ge^* mutants. Subsequently, both the calyptra and pseudoperianth developmentally arrest after between 1 to 2 weeks post fertilization in Mp*knox1-6^ge^* mutants.

**Figure 4.**
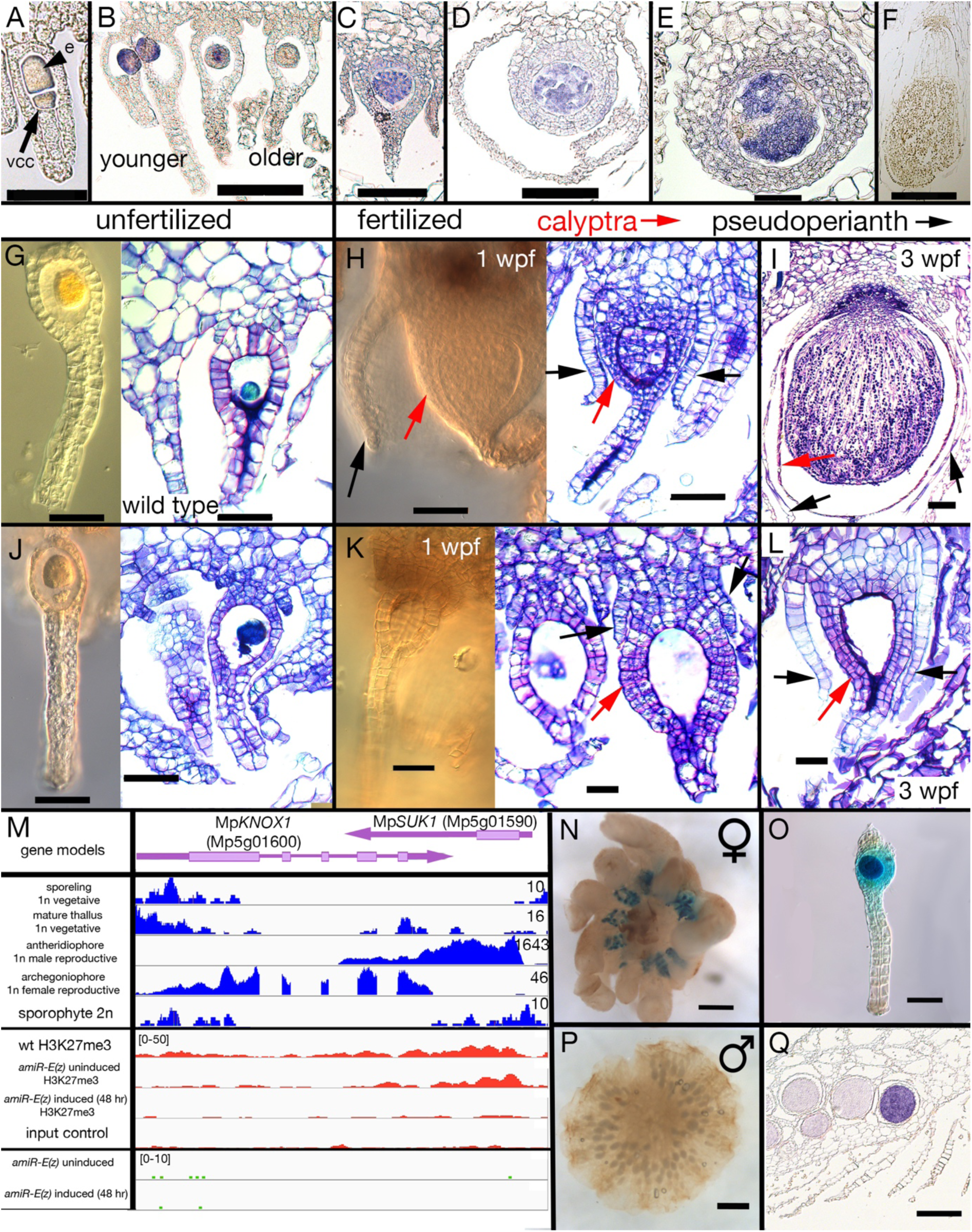
Maternal Mp*KNOX1* is required for post-zygotic embryo development. (A-F) Mp*KNOX1* expression pattern. Mp*KNOX1* transcripts could be detected via *in situ* hybridization in the egg (e) cells of unfertilized archegonia (B), but not in the venter canal cell (A; vcc). After fertilization, expression is detected in young developing sporophytes (approximately 1 week post fertilization, C-D), with expression continuing foot and seta begin to differentiate (E). Mp*KNOX1* is not expressed in older sporophytes where sporogenous tissue has differentiated (F). (**G-L**) Comparison of wild-type and Mp*knox1* development; unfertilized (G, J), 1 week post-fertilization (H, K), and 3 weeks post-fertilization (I, L). Mp*knox1-6^ge^* egg cells appear wild-type-like (unfertilized, G, J). In contrast to wild type (H-I), the embryo of Mp*knox1-6^ge^* mutants does not develop post fertilization (K-L). The zygote fails to undergo cytokinesis, with the nucleus disappearing after three weeks, leaving an empty space (approximately 3 weeks post-fertilization, L). The initial outgrowth of the pseudoperianth (black arrows) and calyptra (red arrows) post-fertilization is not affected in Mp*knox1-6^ge^* mutants, but their development arrests as well approximately 1 week post-fertilization (K-L). (**M**) Based on RNA-seq data and associated gene models, the 3’ end of Mp*KNOX1* overlaps with the 3’ UTR of Mp*SUK1*, which is transcribed from the opposite strand. Note that the RNA-seq data presented in blue have different scales for the different tissues. Predicted full-length Mp*KNOX1* transcripts, i.e. those including exons 2 and 3, are present primarily in the archegoniophore, consistent with the *in situ* data and sqRT-PCR data in Figure 1— figure supplement 1. Mp*KNOX1* and Mp*SUK1* are regulated quite differently. Mp*KNOX1* is neither marked by H3K27me3 in the vegetative gametophyte nor upregulated in the *amiR-*Mp*E(z)* background. In contrast, Mp*SUK1* is expressed at high levels in antheridiophores and is marked by H3K27me3 in the vegetative gametophyte but not upregulated in the *amiR-*Mp*E(z)* background. (**N-P**) Expression of *_pro_*Mp*KNOX1:GUS:SUK1* in an archegoniophore (N), an archegonium (O), and an antheridiophore (P). (**Q**) Signal of Mp*SUK1* in situ hybridization is detected in antheridia of the antheridiophore. Bar = 50 µm (A, E. G, J, K, L, O); 100 µm (B, C, D, H, I); 200 µm (Q); 500 µm (F, N); 1000 µm (P).

At the Mp*KNOX1* locus is a convergently transcribed gene, *SUPPRESSOR OF KNOX1* (Mp*SUK1*, Mp5g01590), which is expressed predominantly in antheridia (**Figure 4M, Q**), suggesting that Mp*KNOX1* may be regulated by an antisense transcript as has been described for Mp*FGMYB* (Mp1g17210) (Hisanaga et al., 2019). Consistent with this hypothesis, a transcriptional fusion of 4.6 kb 5’ of Mp*KNOX1* coding sequence with a GUS reporter coding sequence and 3’ terminator results in expression in both egg cells and antheridia (**Figure 4— figure supplement 1**). In contrast, incorporation 2.7 kb of the Mp*SUK1* antisense transcriptional unit immediately downstream of the GUS coding sequence (*_pro_*Mp*KNOX1:GUS:SUK1*) results in female specific expression (**Figures 4N-P; Figure 4— figure supplement 1)**. However, if a 3’ terminator sequence is inserted between the GUS coding sequence and the Mp*SUK1* sequences the antisense repression is lost, suggesting a requirement of transcriptional overlap but in a non-sequence-specific mechanism (**Figure 4— figure supplement 1**). In contrast to Mp*KNOX1*, Mp*SUK1* is marked with H3K27me3, but its expression was not induced in the *amiR-*Mp*E(z)1* background (**Figure 4M**). Mp*SUK1* also encodes a 225 aa protein that does not overlap with the Mp*KNOX1* coding sequence, but that includes a 55 aa domain conserved across most liverworts.

In addition to the full length Mp*KNOX1* gene, two additional KNOX1-related genes, Mp*KNOX1A* and Mp*KNOX1B*, are encoded in the *M. polymorpha* genome. Both genes encode a conserved KNOX1 MEINOX domain (**Figure 4—figure supplement 2A**), but lack a HD, are predominantly expressed in the antheridia and in sperm cells (**Figure 4—figure supplement 2C**). The three primarily gametophyte-expressed KNOX1 genes were neither marked with H3K27me3 nor induced in the *amiR-*Mp*E(z)1* background (**Figure 4; Figure 4— figure supplement 2B**).

### Mp*BELL234* are highly expressed in antheridia

The acropetal development of antheridia within antheridiophores results in the youngest being peripheral. Mp*BELL2*, Mp*BELL3* and Mp*BELL4* expression was detected in stage 2-4 antheridia (Higo et al., 2016) within antheridiophores as assayed by *in situ* hybridization (**Figure 5A, C, E**). Likewise, translational Mp*BELL2* and transcriptional Mp*BELL3* and Mp*BELL4* β-glucuronidase reporter gene fusion lines exhibit signals in antheridia of developing antheridiophores (**Figure 5B, D, F**). No signal was detected in unfertilized archegoniophores or developing sporophytes in any of the reporter lines, nor was any expression observed in the vegetative gametophyte. However, in contrast to reporter lines, strand-specific transcriptomic analysis suggests that alternative shorter transcripts, likely produced from an alternative promoter, are generated from Mp*BELL3* and Mp*BELL4* in archegoniophores and sporophytes (**Figure 5—figure supplement 1B**). Directed rtPCR experiments confirmed the existence of a short transcript lacking homeodomain sequences produced from the Mp*BELL4* locus during sporophyte development and transcripts encoding homeodomain sequences produced from Mp*BELL3* in archegoniophores and during sporophyte development (**Figure 5—figure supplement 1E**). While sporophyte expression for both genes was observed via *in situ* hybridization (**Figure 5—figure supplement 1E**), potential archegonial Mp*BELL3* expression was difficult to distinguish from background.

**Figure 5.**
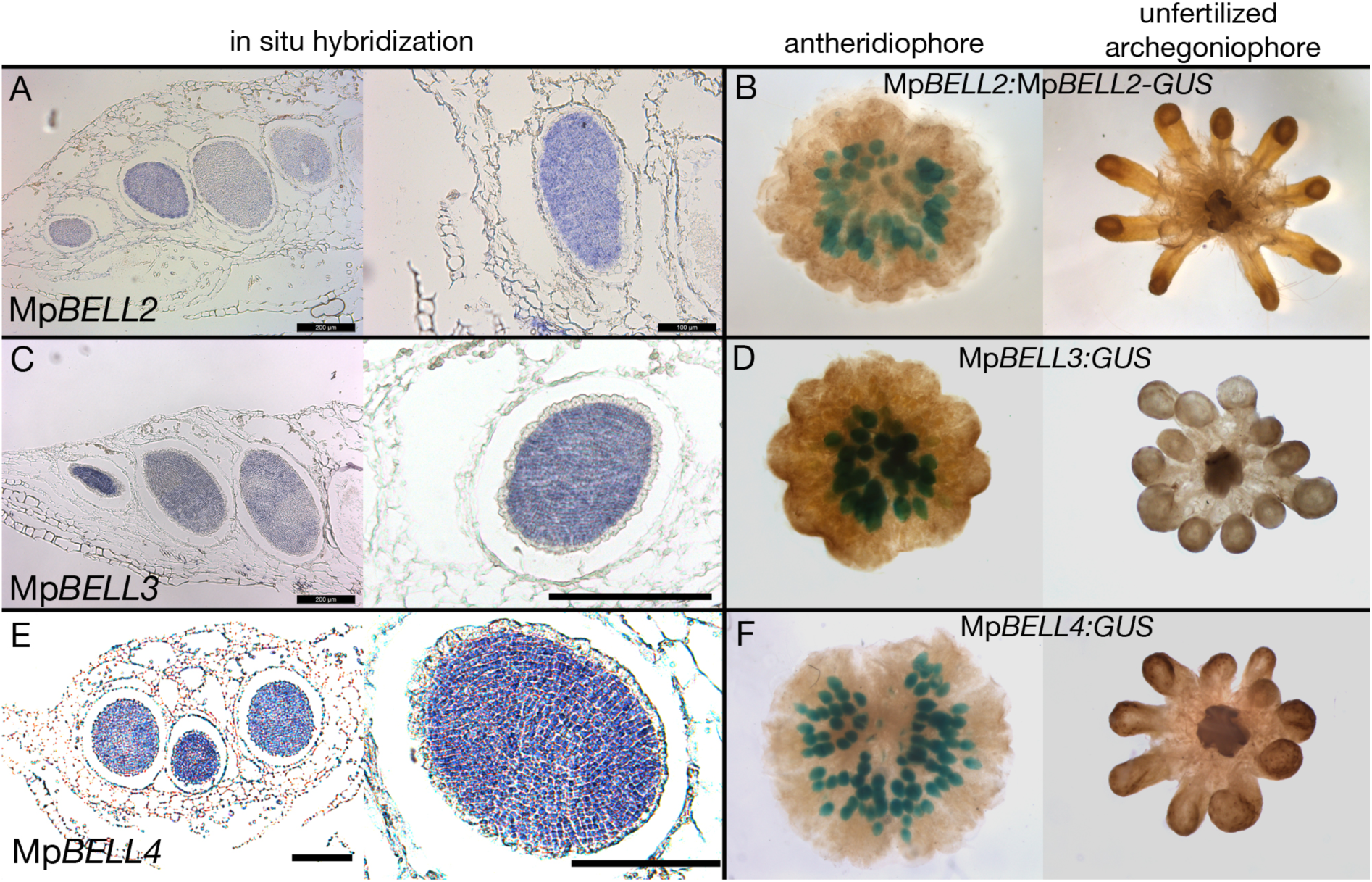
Expression patterns of Mp*BELL2*, Mp*BELL3* and Mp*BELL4* in antheridiophores. *In situ* localisation of all three Mp*BELL* mRNA in young and medium aged antheridia as seen in antheridiophore cross-sections (A, C, E). Antheridiophores of translational (B) and transcriptional (D, F) β-glucuronidase reporter gene fusions of Mp*BELL2* (B), Mp*BELL3* (D) and Mp*BELL4* (F). Signal appears strongest in young to medium aged antheridia [stage 3 and stage 4, (Higo et al., 2016)], with signal being lost in older antheridia towards the centre of the antheridiophore (B, F), possibly due to draining of spermatogenous tissue. Staining was not observed in unfertilized archegonia. Scale bars = 200μm for left panels in A, C, E and right panel in C; 100μm for right panels in A and E.

Of the three primarily antheridiophore-expressed BELL genes, Mp*BELL2* was neither marked with H3K27me3 nor induced in the *amiR-*Mp*E(z)1* background. In contrast, Mp*BELL3* and Mp*BELL4*, experienced a loss of H3K27me3 marks, but were not induced in the *amiR-* Mp*E(z)1* background (**Figure 5—figure supplement 1C-D**). Mp*BELL5* was neither marked with H3K27me3 nor induced in the *amiR-*Mp*E(z)1* background (**Figure 5—figure supplement 1C-D**).

### Paternal MpBELL is required for proper embryo development

To examine whether these MpBELL genes could provide the male counterpart to the female MpKNOX1, we created loss-of-function alleles for each gene (**Table 1**). When used as male parents in crosses with wild-type females, Mp*bell2*, Mp*bell3* and Mp*bell4* single mutants did not exhibit an aberrant phenotype, however, when Mp*bell3* Mp*bell4* (Mp*bell34*) double mutant or Mp*bell2* Mp*bell3* Mp*bell4* (Mp*bell234*) triple mutant males were used in crosses with wild-type females no mature sporophytes were formed. Most fertilization events resulted in sporophyte development that progressed to a globular multicellular stage characteristic of the first week of wild-type development, followed by sporophyte arrest (**Figure 6**). Only occasionally did sporophytes fail to progress beyond zygote formation, resembling the phenotype observed in female Mp*knox1-6^ge^* mutants. In crosses in which mature sporophytes form, the corresponding archegoniophores remain viable until after the sporophytes have matured, while unfertilized archegoniophores undergo senescence. The archegoniophores with arrested sporophytes senesced in manner similar to unfertilized archegoniophores. In contrast, in the reciprocal cross between Mp*bell34* females and wild-type males normal sporophyte development was observed (**Figure 6—figure supplement 1**).

**Figure 6.**
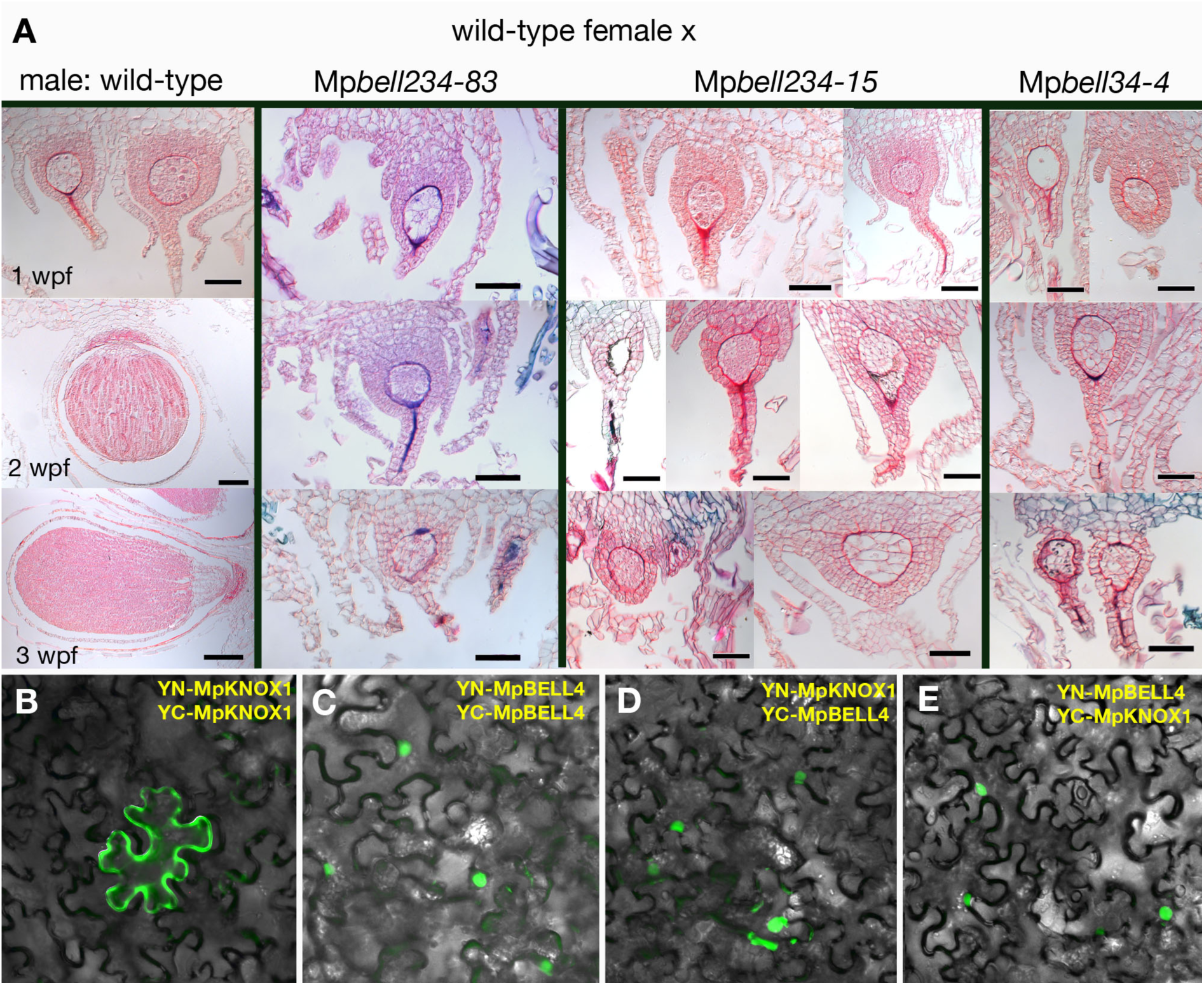
Sporophyte development requires BELL activity in the male gamete. (A) wild-type females crossed with wild-type males (column one) and different Mp*bell* mutant alleles observed after one-, two- and three-weeks post fertilization. Mp*bell234-83^ge^*, Mp*bell234-15^ge^* and Mp*bell34-4^ge^* all show arrested development approximately one week after fertilization. See Table 1 for details on mutant alleles. **(B)** BiFC assay of protein-protein interaction of MpKNOX1 and MpBELL4 in *N. benthamiana* leaves. Homodimers of MpKNOX1 show cytoplasmic localisation, while MpBELL4 homodimers localize to the nucleus. Co-expression of both proteins result in nuclear localisation of heterodimers. Scales bars = 50 µm, except wild-type 2 wpf (100 µm) and wild-type 3 wpf (200 µm).

We next examined whether egg-expressed MpKNOX1 and antheridial-expressed MpBELL proteins could interact. We chose MpBELL4 for analysis due to the truncated nature of MpBELL2 and the extreme length of MpBELL3. In a split YFP BiFC assay, MpKNOX1 on its own is cytoplasmically localized, but when co-expressed with nuclear localized MpBELL4, MpKNOX1 signal becomes nuclear (**Figure 6**). A similar interaction was observed with MpKNOX2 and MpBELL1 (**Figure 6—figure supplement 2**). These interactions are selective, since neither interaction between MpKNOX1 and MpBELL1 nor between MpKNOX2 and MpBELL4 were observed (**Figure 6—figure supplement 2**).

Expression of the fifth *M. polymorpha* MpBELL gene, Mp*BELL5*, may be restricted to the archegoniophore among tissues examined (**Figure 3—figure supplement 3**). As with Mp*KNOX1* (Figure 4) and Mp*FGMYB* (Hisanaga et al., 2019), the Mp*BELL5* transcript may also be opposed by a convergent antheridial-expressed transcript (**Figure 3—figure supplement 2**).

### Activation of sporophyte gene expression in the vegetative gametophyte

The expression patterns of Mp*BELL2/3/4* and Mp*KNOX1* genes are consistent with a role in activating diploid gene expression, and thus we examined whether ectopic co-expression of these genes in vegetative gametophyte is sufficient to activate diploid gene expression (**Figure 6—figure supplement 3A**). Co-expression Mp*KNOX1* and Mp*BELL3* in the vegetative gametophyte is sufficient to activate Mp*KNOX2*, whose expression is normally limited to the sporophyte, and Mp*BELL1*, whose expression is normally limited to the sporophyte or after continuous far-red light induction (Inoue et al., 2019), after 72 hours (**Figure 6—figure supplement 3B**). However, both these genes were also activated by expressing only Mp*BELL3* in the vegetative gametophyte, and furthermore, Mp*KNOX1* was also induced under these conditions (**Figure 6—figure supplement 3B**).

## Discussion

### An ancestral function for TALE-HD genes in the Viridiplantae

The eukaryotic life cycle alternates between haploid and diploid phases, initiated by meiosis and gamete fusion, respectively. Organisms spanning the phylogenetic diversity of eukaryotes, including ascomycete and basiomycete fungi, Amoebozoa, brown algae, and Chlorophyte algae, have also been shown utilize paralogous homeodomain proteins that heterodimerize following gamete fusion to initiate the diploid genetic program, lending support the idea that this may have been the ancestral function of homeodomain proteins. Our observations in *M. polymorpha* indicate that TALE-HD proteins, specifically MpKNOX1 and MpBELL2/3/4, are initially supplied in gametes (egg and sperm, respectively), and that this gametic expression is required for diploid sporophyte development (**Figure 7**). These data are consistent with a general model for the Viridiplantae whereby KNOX is supplied via the egg (equivalent to the (-) gamete in *Chlamydomonas*) and BELL via the sperm (equivalent to the (+) gamete in *Chlamydomonas*), and that once together in the zygote, the diploid genetic program can be activated. Thus, the basic tenants of the genetic regulation of the haploid to diploid transition elucidated in another lineage of Viridiplantae also apply to a basal lineage of land plants, consistent with the notion that such a system was present in the common ancestor of extant Viridiplantae.

**Figure 7.**
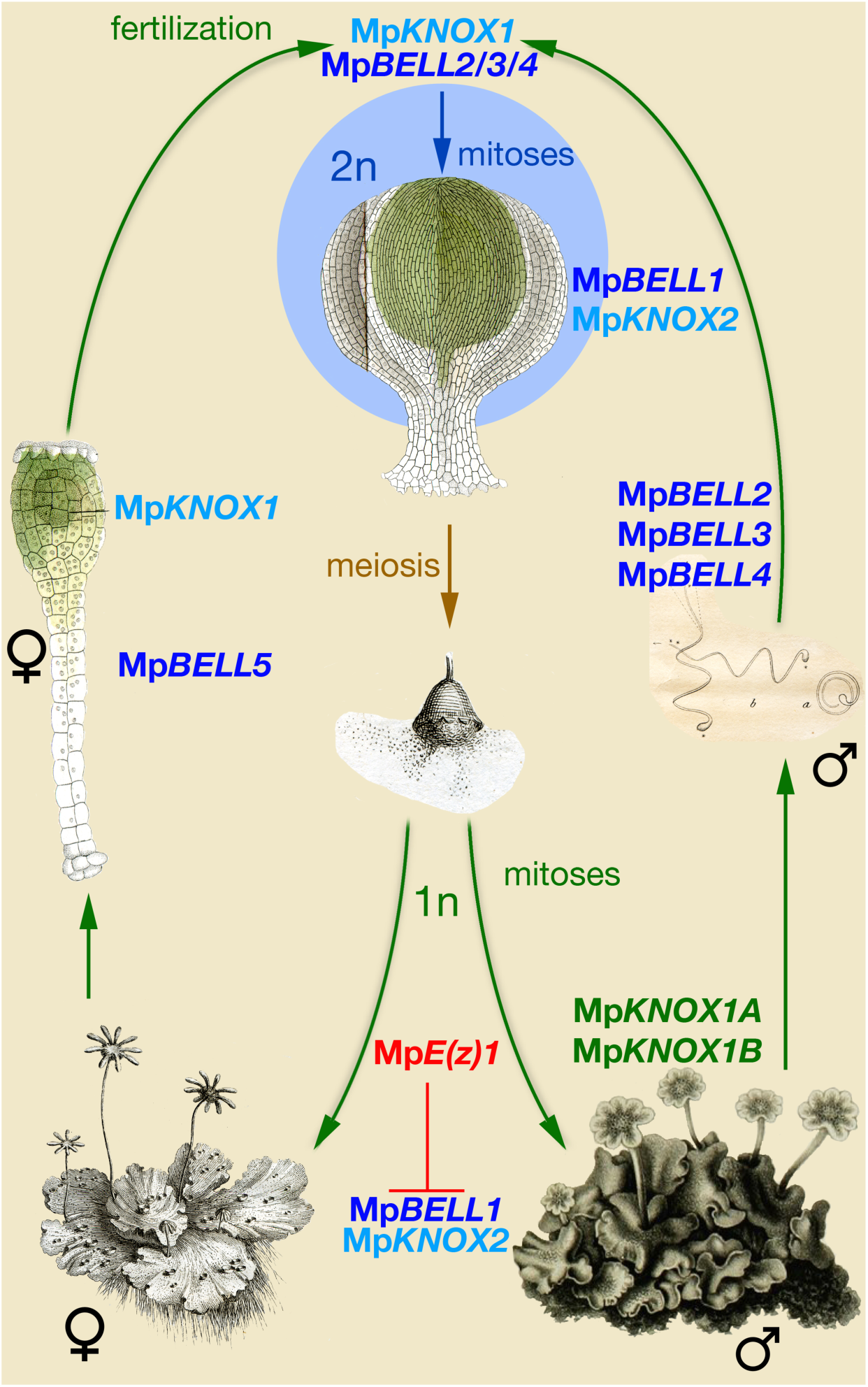
Model of KNOX and BELL function in *Marchantia polymorpha*. Three Mp*BELL* genes are expressed in antheridia, while a single Mp*KNOX1* is expressed in the egg cell of the archegonia. After fertilization, MpBELL2/3/4 and MpKNOX1 proteins interact to activate the zygotic program. Mp*BELL1* and Mp*KNOX2* are expressed at later stages of the sporophyte development. In the haploid phase of the life cycle, Mp*E(z)1* represses sporophyte-expressed Mp*BELL1* and Mp*KNOX2*. The homeo-domain lacking Mp*KNOX1A* and Mp*KNOX1B* are expressed with antheridia specific Mp*BELL2/3/4*, potentially to prevent functionality by sequestering MpBELL proteins. The function of Mp*BELL5* is unknown. Images from (Marchant, 1713, Mirbel, 1835, Thuret, 1851, Unger, 1837).

While crosses with Mp*knox1* females always resulted in zygotic arrest, this phenotype was seldom seen in crosses involving Mp*bell234* males, but rather, in the latter case a limited amount of embryo development ensued before developmental arrest. Note that this is the opposite of what might have been expected based on ectopic expression of these genes in the vegetative gametophyte where Mp*BELL3* alone was sufficient to activate some diploid gene expression whereas Mp*KNOX1* was not (**Figure 6—figure supplement 3**). There exist several possible, not necessarily mutually exclusive, explanations for the discordance between these two reciprocal crosses. First, the Mp*bell* alleles may not be null — despite large internal deletions each still encodes an intact conserved carboxyl homeodomain (**Figure 1**), and perhaps a truncated MpBELL protein is sufficient for limited activity. A corollary of this possibility is that maternal transcripts at the Mp*BELL3* locus (**Figure 5—figure supplement 1**) could contribute to initiating embryo development, but are not sufficient for its continuation beyond a rudimentary ball of cells. Third, embryo gene expression could be induced independently of paternally supplied Mp*BELL34*, perhaps via upstream regulators of Mp*BELL34* being supplied to the zygote by the sperm. This maternal zygotically produced MpBELL34 could then heterodimerize with MpKNOX1 and initiate the embryo genetic program. Such a scenario is reminiscent of *Chlamydomonas* where, in the absence of GSM1 (KNOX), GSP1 (BELL) can induce diploid gene expression (Kariyawasam et al., 2019). It was proposed that in wt(+) x *gsm1*(-) zygotes GSM1 may be activated by its upstream regulator MID supplied by the (-) gamete (Kariyawasam et al., 2019). Fourth, maternal expression of Mp*BELL5* could play a role, perhaps by forming heterodimers, with MpKNOX1 upon a fertilization stimulus, able to initiate, but not sustain, embryo development.

Given the phylogenetic affinity of mosses as a sister group to liverworts (Renzaglia et al., 2018), we suggest that expression of Pp*BELL1* in the sperm as observed by Ortiz-Ramírez et al. (Ortiz-Ramírez et al., 2017) is genuine and that the earlier reported lack of male expression (Horst et al., 2016) may be related to sterility in the particular *Physcomitrella* strain (Meyberg et al., 2019). The phenotype of Mp*knox1* alleles is also more extreme than null loss-of-function alleles of all three PpKNOX1 genes where a sporophyte with viable spores is still formed (Sakakibara et al., 2008). Hence, aspects of the ancestral system regulating the haploid to diploid transition could be present in multiple early diverging land plant lineages, albeit with *M. polymorpha* retaining more ancestral attributes than *Physcomitrella*. In contrast, in angiosperms it appears paternally supplied pluripotency factors related to BABYBOOM, an AP2/ERF transcription factor, can activate the zygotic genetic program (Khanday et al., 2019, Conner et al., 2017, Conner et al., 2015). Furthermore, despite the extensive literature on the functions of KNOX/BELL in angiosperms, the only potential known vestige of this ancestral system existing in angiosperms is the presence of KNOX2 function in the Arabidopsis female gametophyte, possibly in the egg cell (Pagnussat et al., 2007). Thus, it seems that the ancestral function has been entirely lost in some derived land plants, as it has presumably in the metazoan lineage (Bowman et al., 2016b).

Since liverworts are placental organisms, with the diploid sporophyte nourished by the maternal haploid gametophyte, there is opportunity for extensive intergenerational communication. For example, that the maternal gametophytic calyptra and pseudoperianth initiate their development in Mp*knox1* mutants implicates a non-cell autonomous zygote derived signal activated post-fertilization in an Mp*knox1* independent manner. Likewise, the premature senescence of archegoniophores fertilized by Mp*bell34* males and bearing aborted embryos suggests a signal emanating from older sporophytes to maintain the viability of the maternal archegoniophore tissues until sporophyte development is completed.

### Roles of MpKNOX1 in the Marchantia sporophyte

One of the key innovations of land plants was the evolution of a mutlicellular diploid generation, the embryo, via mitoses interpolated between gamete fusion and meiosis (Bower, 1908). In seed plants, ferns, and mosses KNOX1 activity is associated with continued cell proliferation, including sporophyte apical meristem activity (Hay and Tsiantis, 2010, Sakakibara et al., 2008, Sano et al., 2005). While *M. polymorpha* and other liverworts lack apical cells and meristems during sporophyte development (Kienitz-Gerloff, 1874), Mp*KNOX1* expression is detected throughout developing sporophytes until the inception of sporogenous cell differentiation, consistent with Mp*KNOX1* having a role in cell proliferation during *M. polymorpha* sporophyte development. Thus, while MpKNOX1 retains the ancestral function in the haploid to diploid transition, the lack of zygotic cell division in Mp*knox1* mutants and the Mp*KNOX1* sporophytic expression pattern suggest that neofunctionalization of KNOX1 contributed to the evolution of the embryo via stimulation of cell proliferation.

In eukaryotic lineages in which multicellularity evolved in the diploid phase of the life cycle, it is perhaps not surprising that genes involved in the haploid to diploid transition, or paralogs thereof, have been co-opted into roles directing development of the diploid generation. In the basidiomycete fungus *Coprinopsis*, the same HD heterodimer that initiates the diploid genetic program also directs early developmental stages in the multicellular diploid (Kamada, 2002). Most conspicuously, despite the loss of their role in zygotic gene activation, both non-TALE (e.g. Hox) and TALE HD genes act to pattern the metazoan body (Merabet and Mann, 2016, Pearson et al., 2005, Lewis, 1978). In the syncytial Chlorophyte alga *Caulerpa lentillifera*, differential expression of TALE-HD genes in the diploid body was speculated to influence differentiation of fronds and stolons, but functional data are lacking (Arimoto et al., 2019). Finally, in land plants, as the multicellular sporophyte evolved increasing complexity, KNOX/BELL genetic modules were co-opted to direct development of novel organs and tissues via regulation of the extent of proliferation (KNOX1) or promotion of differentiation (KNOX2) in conjunction with other gene regulatory networks (Hay and Tsiantis, 2010, Furumizu et al., 2015).

### *Marchantia* gametophyte and sporophyte TALE-HD expression is distinctly regulated

The two TALE-HD genes expressed predominantly in during sporophyte development, Mp*KNOX2* and Mp*BELL1*, are actively repressed in the vegetative gametophyte via Mp*E(z)1,* i.e. marked with H3K27me3, a hallmark of polycomb mediated repression (**Figure 3**). The active suppression of Mp*BELL1* and Mp*KNOX2* in the vegetative gametophyte of *M. polymorpha* resembles the situation in *P. patens* where loss-of-function alleles disrupting PRC2 activity result in ectopic gametophytic PpKNOX1 activity, with sporophyte-like bodies developing in the haploid generation (Mosquna et al., 2009, Okano et al., 2009, Pereman et al., 2016). One intriguing observation was the rapidity with which Mp*KNOX2* and Mp*BELL1* were de-repressed in the inducible *amiR-*Mp*E(z)1* lines, with expression detected as early as two hours post induction. That this happens within a couple hours suggest that that PRC2 mediated repression of Mp*KNOX2* and Mp*BELL1* is both active and labile (Di Croce and Helin, 2013).

Of the three BELL genes (Mp*BELL2/3/4*) expressed specifically in the antheridia during sperm differentiation, Mp*BELL3* and Mp*BELL4* are marked with H3K27me3 in the vegetative gametophyte, but MpB*ELL2* is not. However, none of these three genes are de-repressed in *amiR-*Mp*E(z)1* lines (**Figure 5—figure supplement 1**). Likewise, the three KNOX1-related genes whose expression is predominantly gametophytic are neither marked with H3K27me3 in the vegetative gametophyte nor de-repressed in *amiR-*Mp*E(z)1* lines (**Figure 4, Figure 4—figure supplement 2**). In contrast to Mp*KNOX1*, Mp*SUK1* appears to be marked by H3K27me3 in the vegetative gametophyte, but whether the mark is sex-specific is unknown. Collectively, these data indicate that TALE-HD genes expressed during later sporophyte development are regulated in a fundamentally different manner than the gamete-expressed TALE-HD genes. Further, de-repression of Mp*KNOX2* and Mp*BELL1* in *amiR-*Mp*E(z)1* lines implies that their activators are present in the vegetative gametophyte, while activators of the gamete-expressed TALE-HD genes are not present in this tissue.

### TALE-HD genes with converse expression patterns

The two KNOX1-related sequences lacking homeodomains are reminiscent of similar, albeit independently evolved, proteins in angiosperms that have been demonstrated to act as inhibitors of KNOX/BELL function by forming inactive heterodimers primarily with BELL partners (Magnani and Hake, 2008, Kimura et al., 2008). The gametophytic expression, especially in antheridia and sperm cells, of the homeodomain lacking Mp*KNOX1A* and Mp*KNOX1B* genes prompts the hypothesis that the encoded proteins may act in a similar manner, perhaps as a safeguard against inappropriate activity MpBELL in the male gametophyte. This is particularly pertinent since ectopic expression of Mp*BELL3* alone in the vegetative gametophyte can activate Mp*KNOX1*, Mp*KNOX2* and Mp*BELL1* (**Figure 6—figure supplement 3**). Consistent with a protective role is the lack of aberrant phenotype observed in lines constitutively expressing Mp*KNOX1* in male gametophytes (Dierschke and Bowman, unpublished). In contrast to the antheridial or sporophytic expression of the other MpBELL genes, Mp*BELL5* transcripts are detected in archegoniophores. Given the strong heterodimerization affinity preferences of BELL proteins with KNOX proteins (Bellaoui et al., 2001, Joo et al., 2018), the only conspicuous potential partner in *M. polymorpha* would be MpKNOX1. Hence, Mp*BELL5* might play an as yet undefined role in the activation of zygotic gene expression. At the Mp*BELL5* locus, a putative convergently transcribed gene is expressed predominantly in antheridia (**Figure 5—figure supplement 1**), suggesting Mp*BELL5* may be regulated by an antisense transcript in a manner similar to Mp*KNOX1* (**Figure 4**) and Mp*FGMYB* (Hisanaga et al., 2019). Thus, male-expressed antisense transcript-mediated repression could provide a general mechanism for female-specific expression of autosomal genes whose regulation may be linked to the feminizing-locus on the female sex chromosome (Bowman, 2016, Knapp, 1935, Lorbeer, 1936).

### *Marchantia* BELL gene diversity resembles that of Viridiplantae algae

The phylogenetic diversity of BELL genes in *M. polymorpha* more closely resembles the diversity observed in charophyte and Chlorophyte algae than that of other land plants (Joo et al., 2018, Lee et al., 2008). All previously described land plant BELL genes form a single clade that evolved within the charophycean algae (**Figure 2**). The single *M. polymorpha* gene orthologous to most other land plant genes, Mp*BELL1*, is the only one predominantly expressed during sporophyte development. While there is little phylogenetic resolution of the other BELL genes, their diversity suggests an ancient proliferation of BELL paralogs within the charophycean algal lineage leading to land plants. *M. polymorpha* retained this diversity that was subsequently lost in some derived land plant lineages. The *Chlamydomonas* genome encodes two BELL paralogs, *GSP1* and *HDG1*. *HDG1* is expressed in both (+) and (-) gametes suggesting a role in reproduction, but the function of which is unknown. Likewise, the expression pattern of Mp*BELL5*, a member of the GLX-basal clade that evolved early in the charophytes, also suggests an as yet unresolved function in reproduction. Finally, the structural (two or three homeodomains) and extensive sequence diversity of the Mp*BELL2/3/4* paralogs is consistent with the rapid evolution of genes involved reproductive processes (Swanson and Vacquier, 2002).

### An ancestral function for TALE-HD genes in eukaryotes

In the Viridiplantae (*Chlamydomonas* and *Marchantia*) the haploid to diploid transition is mediated by two TALE-HD genes. In the red alga *Pyropia yezoensis* KNOX gene expression is detected in the diploid conchosporangium, but not in haploid thalli (Mikami et al., 2019). Since *Pyropia*, *Chlamydomonas* and *Marchantia* span much of the phylogenetic diversity of the Archaeplastida, the common ancestor of this group likely utilized TALE-HD proteins of the KNOX and BELL subfamilies mediate the haploid to diploid transition. The phylogenetic distribution of KNOX and BELL subfamilies and their heterodimerization affinities (Joo et al., 2018), suggests this can likely be extended to all Archaeplastida. In contrast, in both ascomycete and basidiomycete fungi the haploid to diploid transition is mediated by heterodimerzation of a TALE-HD and a non-TALE HD protein (Gillissen et al., 1992, Goutte and Johnson, 1988, Herskowitz, 1989, Hull et al., 2005, Kues et al., 1992, Spit et al., 1998, Urban et al., 1996). In the Amoebozoa *Dictyostelium*, the two homeodomain-like proteins controlling the haploid to diploid transition are highly divergent, rendering the phylogenetic affinities enigmatic (Hedgethorne et al., 2017). Finally, in the brown alga *Ectocarpus* two TALE-HD proteins, OUROBOROS and SAMSARA, mediate the transition, however, their lack of gamete specific expression raises questions concerning the precise mechanism (Arun et al., 2019). As these taxa span eukaryotic phylogenetic diversity, one role of homeodomain genes in the ancestral eukaryote was to regulate the haploid to diploid transition — the evolution of the homeodomain in the ancestral eukaryote was associated with evolution of a novel life cycle (**Figure 7—figure supplement 1**). In arguably the most intensively studied taxon, the Metazoa, zygotic gene activation has been replaced by maternally derived pluripotency factors (Schulz and Harrison, 2019), but homeodomain genes have been retained.

Given the ancestral eukaryote possessed a minimum of two HD genes, including one TALE-HD and one non-TALE-HD (Bharathan et al., 1997, Burglin, 1997, Derelle et al., 2007), it is an intriguing question as to whether, ancestrally, both TALE and non-TALE paralogs acted in the diploid to haploid transition. The two possible ancestral scenarios (TALE + non-TALE or TALE + TALE), require either a co-option of a pre-existing paralog (e.g. non-TALE), or alternatively, a TALE gene duplication and transference of function to the new paralog, respectively. To resolve the ancestral condition broader phylogenetic sampling and functional analyses across additional unicellular Bikont eukaryotic lineages, particularly in the Excavata, and the paraphyletic sister groups of the Metazoa (Choanoflagellates, Filasterea, and Ichthyosporea) might be informative (**Figure 7—figure supplement 1**). Resolution of the ancestral condition could inform whether primary ancestral function of both TALE-HD and non-TALE-HD was regulating the haploid to diploid transition or whether the non-TALE-HD genes had another fundamental function in ancestral eukaryote.

## Acknowledgements

We thank our colleagues at the Joint Genome Institute (JGI); the work conducted by the U.S. Department of Energy (DOE) JGI, a DOE Office of Science User Facility, is supported by the Office of Science of the U.S. DOE under contract DE-AC02-05CH11231. This work was supported by Monash University and funding from the Australian Research Council (FF0551326, DP130100177; DP170100049) to J.L.B. S.Z. acknowledges support from the German Research Foundation (SFB944, P13). We thank Keisuke Inoue for providing pMpGE_En04, pBC-12, pBC-14, pBC-23, pBC-34 and Edwin Lampugnani for BiFC vectors. We also thank Claudia Gieshoidt for *Marchantia* tissue culture support.

## Materials and Methods

### Artificial microRNA design

We used the same amiR as described in Flores-Sandoval *et al*. (Flores-Sandoval et al., 2016).

### Ecotypes, plant growth, transformation and induction

*M. polymorpha* ssp *ruderalis*, ecotype BoGa obtained from the Botanical Garden of Osnabrueck, Germany was used in SZ lab. Plant cultivation, transformation and induction was according to Ishizaki et al. (Ishizaki et al., 2008) and Althoff et al. (Althoff et al., 2014). *M. polymorpha* ssp *ruderalis*, ecotype MEL was used in JLB lab and grown, transformed and induced according to Ishizaki et al. (Ishizaki et al., 2008) and Flores-Sandoval et al. (Flores-Sandoval et al., 2015). In case of double transformations, sporelings were co-transformed using two constructs featuring different selectable markers. Spores were drained and plated on selection media for approximately 2 weeks and then subjected to a second round of selection before being transferred to ½ B5 plates. For induction of reproductive organs, plants were transferred to white light supplemented with far red light (735 nm; 45 lmol/m2/s) on ½ B5 media supplemented with 1 % glucose.

### Genotyping, RNA extraction, cDNA synthesis and sqRT-PCR

DNA was extracted using a modified protocol from Edwards *et al*. (Edwards et al., 1991). Instead of vacuum drying, the pelleted DNA was air-dried. Amplicons were directly sequenced. RNA was extracted from ≈100mg wet weight tissue using RNeasy kit from Qiagen following the manufacturers instruction including an on-column DNase I digestion. The same amount of total RNA was used for cDNA synthesis using either Superscript II (Invitrogen) [SZ lab] or Bioscript (Bioline) [JLB lab] reverse transcriptase according to the manufacturer’s instructions with oligo-dT_15_ primer.

**Table 2.**
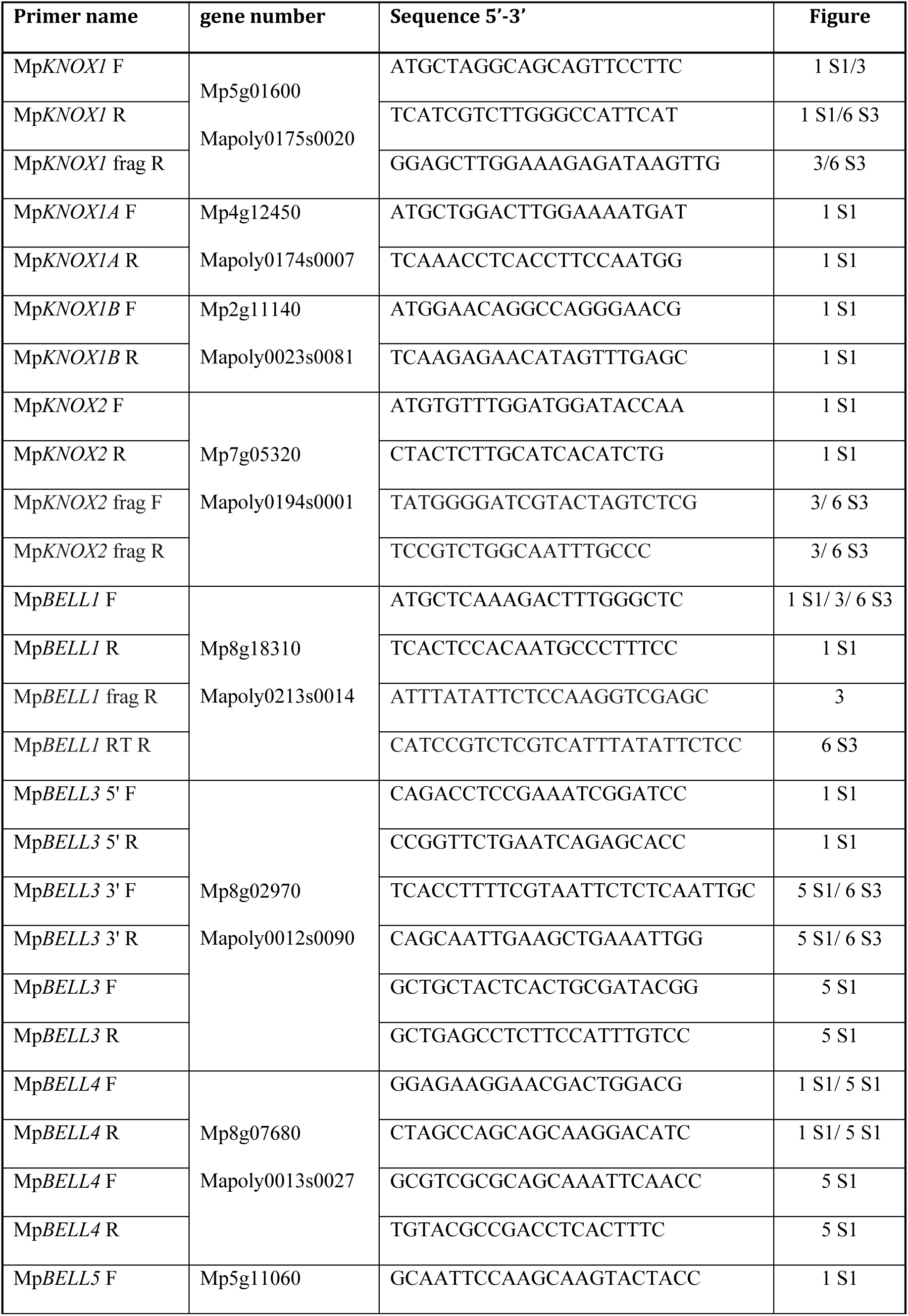
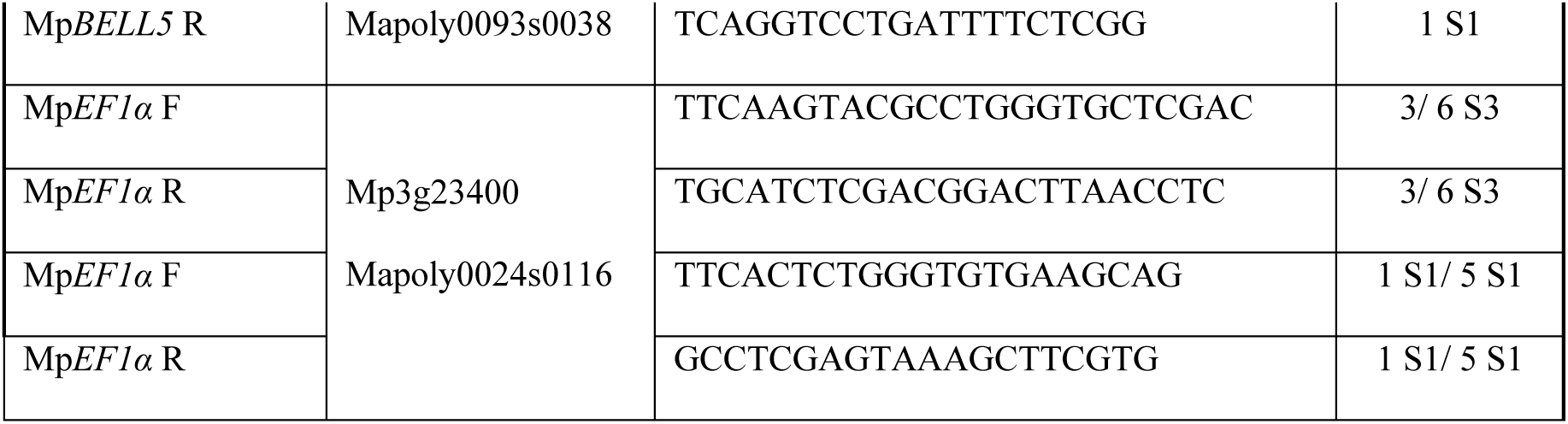
Primer name, gene number, sequence and corresponding Figure used for sqRT-PCR

### In situ hybridisation, sectioning, staining and microscopy

*M. polymorpha* ssp *ruderalis*, ecotype BoGa tissue fixation, embedding, sectioning and hybridization with DIG-labelled antisense RNA probes were performed according to Zachgo (Zachgo, 2002). Sections were either stained in toluidine blue O (TBO) alone or counterstained with ruthenium red (RR) according to Retamales and Scharaschkin (Retamales and Scharaschkin, 2014). Staining time for RR and TBO was 90 sec and 60 sec, respectively. Microscopic slides were observed using a Leica MZ16 FA microscope and pictures were taken with a Leica DFC490 camera. Plants were observed using either a Lumar dissecting microscope (Zeiss) and photographed with AxioCam HRc and and AxioVision software (both Zeiss) or a Leica stereomicroscope (Leica M165 FC) and photographed with an integrated Leica DFC490 camera.

**Table 3.**
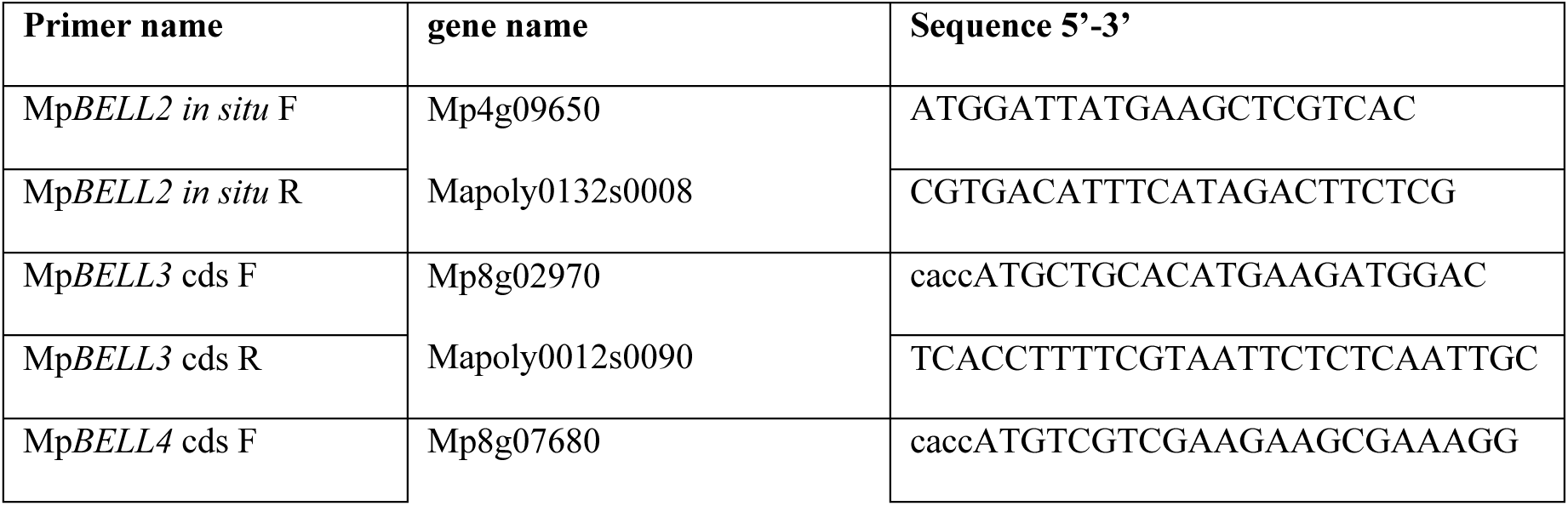
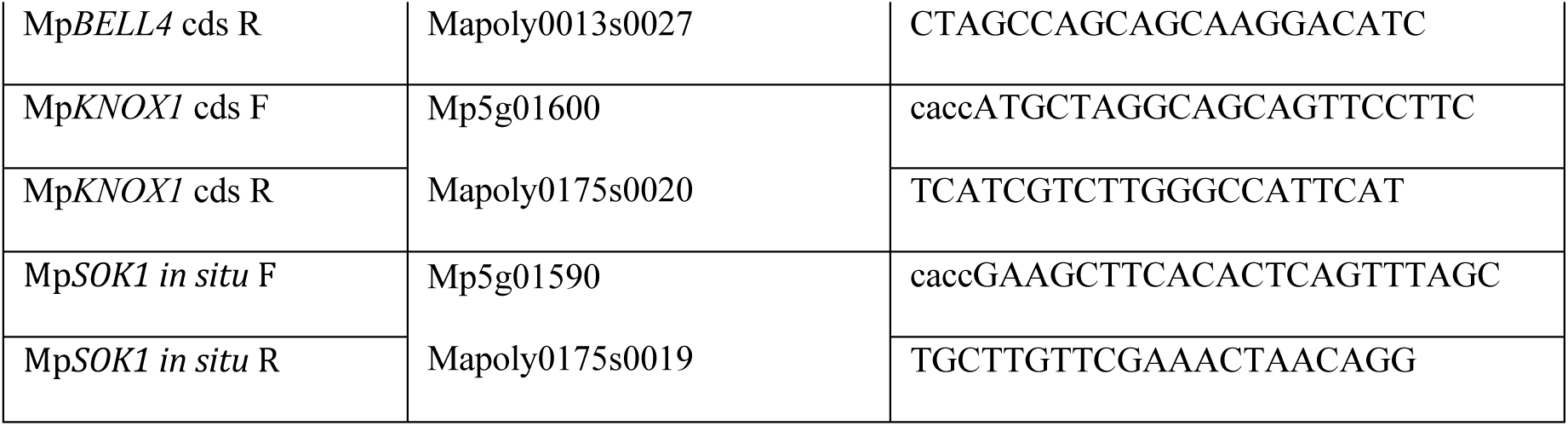
Primer name, gene name and sequence used for *in situ* probe preparation

### Bimolecular fluorescence complementation

Coding sequences of KNOX and BELL genes were seamlessly cloned into the pGREEN derivatives (Hellens et al., 2000) pAMON/pSUR for N-terminal split VENUS (I152L) fusion and into pURIL/pDOX for C-terminal split VENUS (I152L) fusion (Lampugnani et al., 2016). All constructs feature the 35S promoter driving the N- (pAMON/pURIL) or C-terminal (pSUR/pDOX) part of the split VENUS (I152L) fluorophore for translational fusion of the gene-of-interest in frame with the split VENUS (I152L). All constructs were transformed into *A. tumefaciens* strain GV3001 for transfection of plant leaves (Bracha-Drori et al., 2004). For Fluorescence microscopy usually 3, approximately 50 x 50 mm, abaxial leaf tissue fragments were examined under an Axioskop 2 mot plus (Zeiss) microscope and photographed using AxioCam HRc and AxioVision software. For YFP filter set 46 (excitation BP 500/20; beam splitter FT 515; emission BP 535/30 was used.

### Overexpression and transcriptional and translational fusion constructs

Primer used for overexpression with the endogenous, constitutive Mp*EF1α* promoter (Althoff et al., 2014), inducible overexpression and transcriptional and translational fusion constructs are shown below. The complete 1.8kb Mp*KNOX1* coding sequence and a 4.8kb fragment of the Mp*BELL3* coding sequence were amplified from cDNA and seamlessly cloned (NEBuilder, NEB) into pENTR-D (Invitrogen) via *Not*I/*Asc*I sites. The Mp*KNOX1* coding sequence was subsequently recombined into pMpGWB403 (Addgene entry #68668) for constitutive expression, whereas the Mp*BELL3* coding sequence was recombined into the estrogen inducible binary pHART XVE (Flores-Sandoval et al., 2016). Regulatory sequences upstream of the transcriptional start site of Mp*KNOX1* (4.6kb), Mp*BELL3* (5.6kb) and Mp*BELL4* (5.8kb) were amplified and cloned into pENTR-D and subsequently recombined using LR-clonase II (Invitrogen) into pMpGWB104 (Addgene entry #68558) featuring the β-glucuronidase reporter gene. To elucidate the transcriptional regulation at the Mp*KNOX1*/Mp*SUK1* locus, the 4.6kb regulatory region of Mp*KNOX1* was seamlessly cloned (NEBuilder, NEB) into the *Hind*III/*Xba*I site 5’ of the Gateway cassette of pMpGWB401 (Addgene enry #68666). The reversely transcribed Mp*SUK1* locus (2.7kb) was seamlessly cloned into either the *Sac*I site or *Asc*I site (5’ or 3’ of the NOS terminator). Subsequently the β-glucuronidase reporter gene was recombined using pENTR-GUS (Invitrogen). For translational fusions of Mp*BELL2*, Mp*KNOX1A* and Mp*KNOX1B*, the regulatory sequence 5’ upstream of the transcriptional start site including the genomic gene locus excluding the stop codon was amplified and cloned into pRITA, a shuttle vector featuring the β-glucuronidase reporter gene and NOS terminator. Mp*BELL2* and Mp*KNOX1*A were cloned into the *Kpn*I or *Sal*I/*Kpn*I site, respectively. Mp*KNOX1B* was seamlessly cloned using Infusion cloning (Clontech). All constructs were subsequently transferred to the binary vector pHART using *Not*I.

**Table 4.**
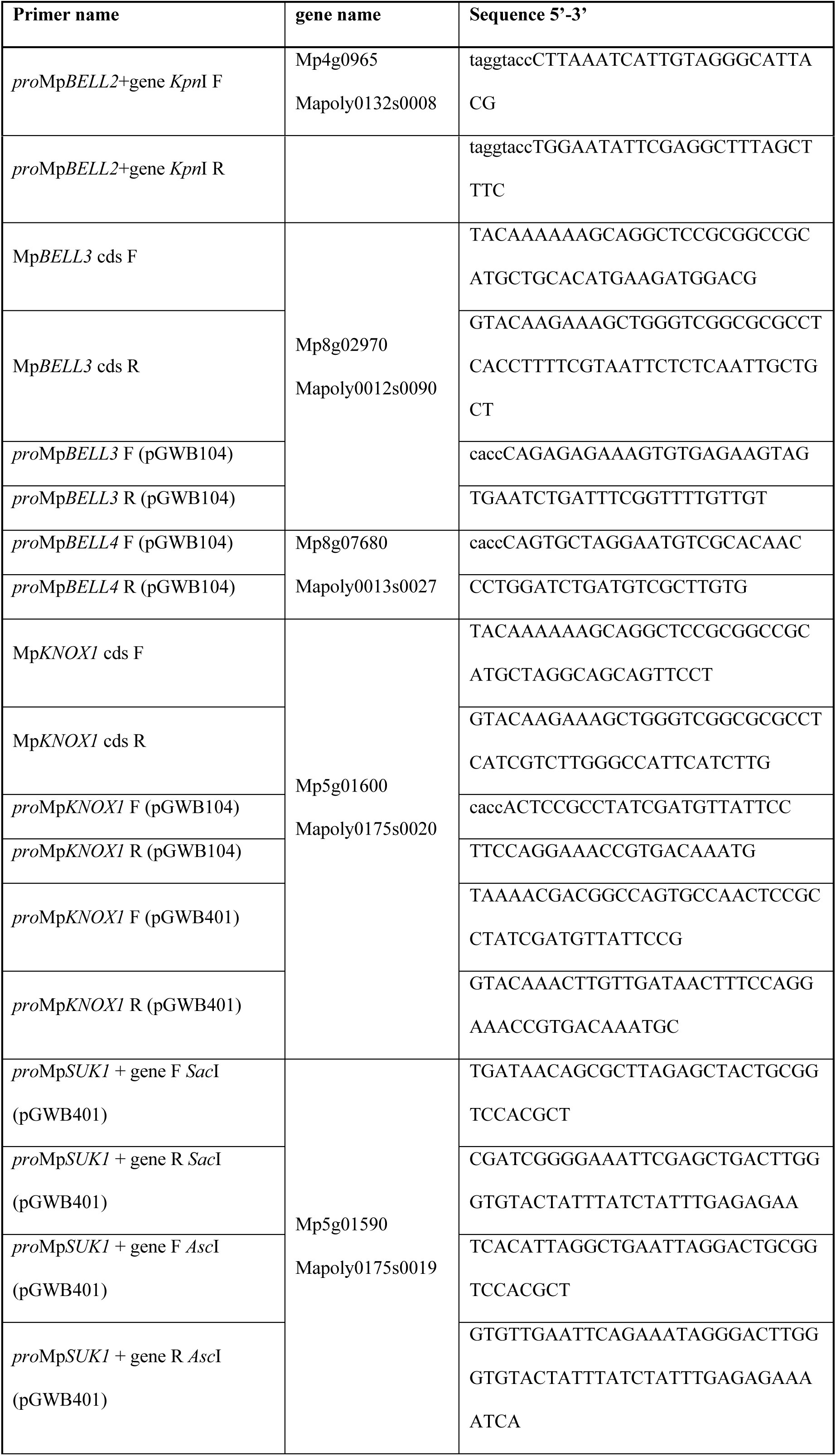
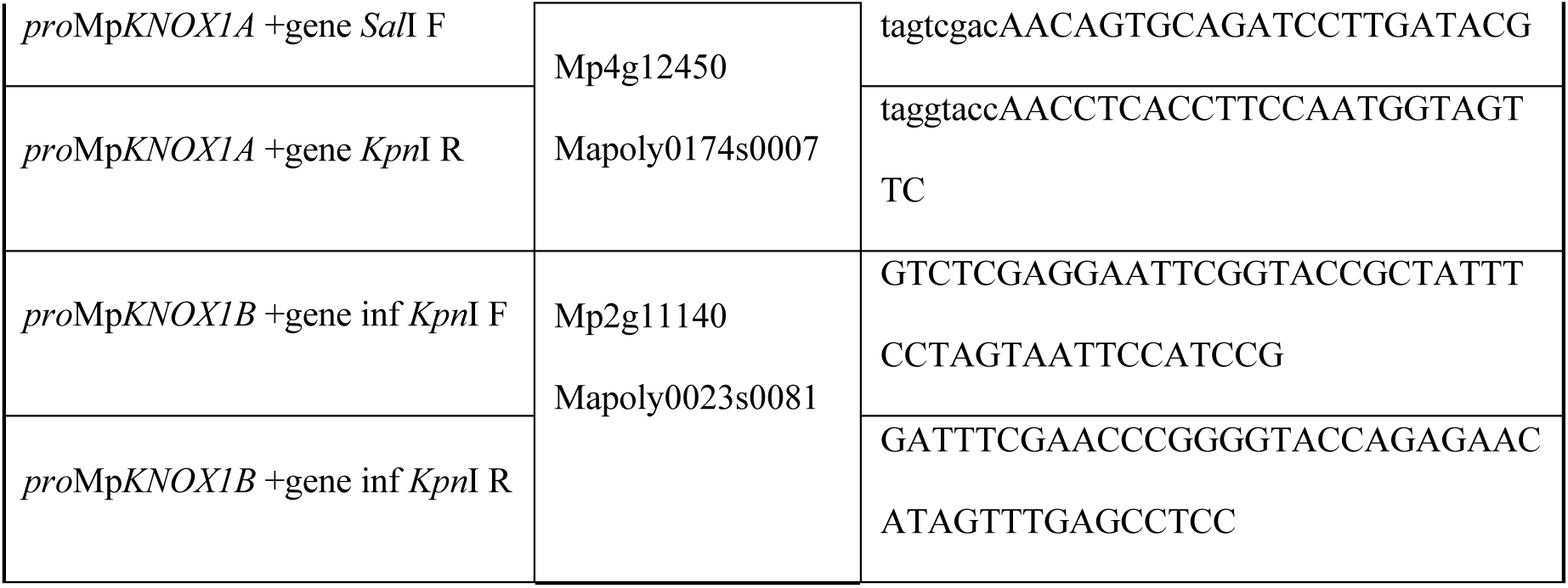
Primer name, gene name and sequence used for overexpression, inducible overexpression, and transcriptional and translational β-glucuronidase reporter constructs

### CRISPR/Cas9 mediated gene knock-out

Potential gRNAs were obtained and screened for off targets in the *M. polymorpha* genome using CRISPOR website (http://crispor.tefor.net/). Quadruple gRNAs were each cloned into pMpGE_En04, pBC-14, pBC-23 or pBC-34 using *Bsa*I sites. Double gRNAs were cloned into pMpGE_En04 and pBC-14 (Hisanaga et al., 2019). These guides were subsequently assembled into the pMpGE_En04 backbone using *Bgl*I sites. pMpGE_En04 were gateway cloned into pMpGE010 [(Sugano et al., 2018, Sugano et al., 2014, Hisanaga et al., 2019); Addgene entry #71536] featuring Cas9 nuclease. In case of 6 gRNAs, the remaining 2 were cloned into pMpGWB401 [(Ishizaki et al., 2015); Addgene entry #68666] and plants doubly transformed (Ishizaki et al., 2008).

**Table 5.**
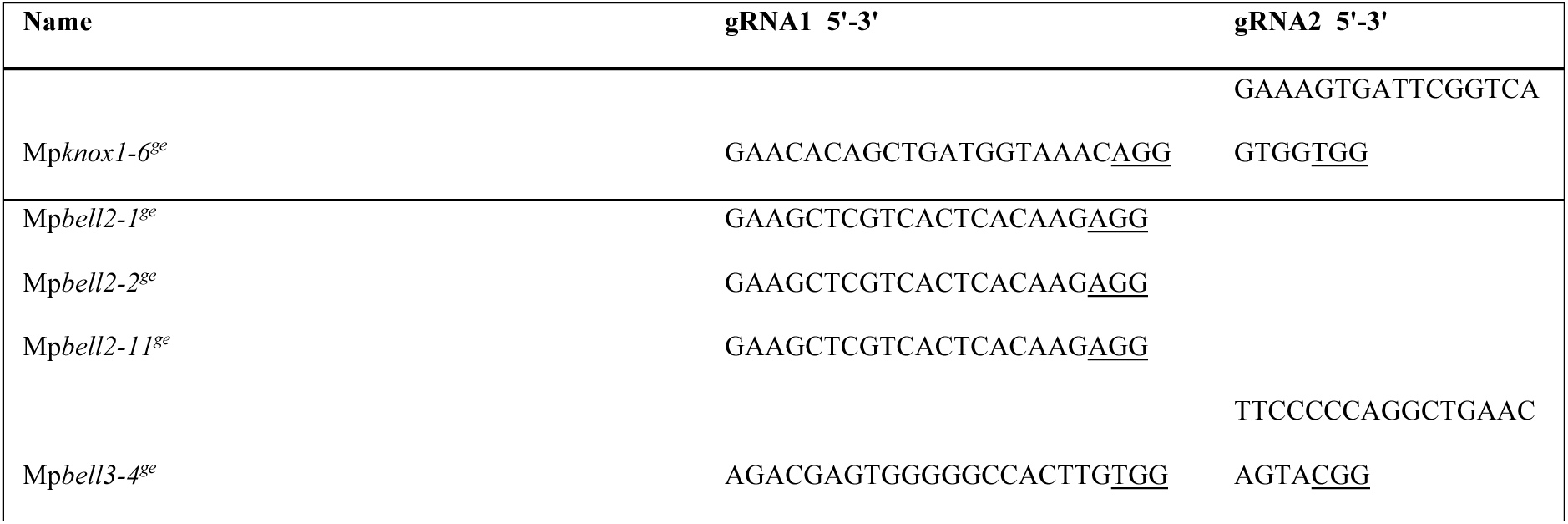
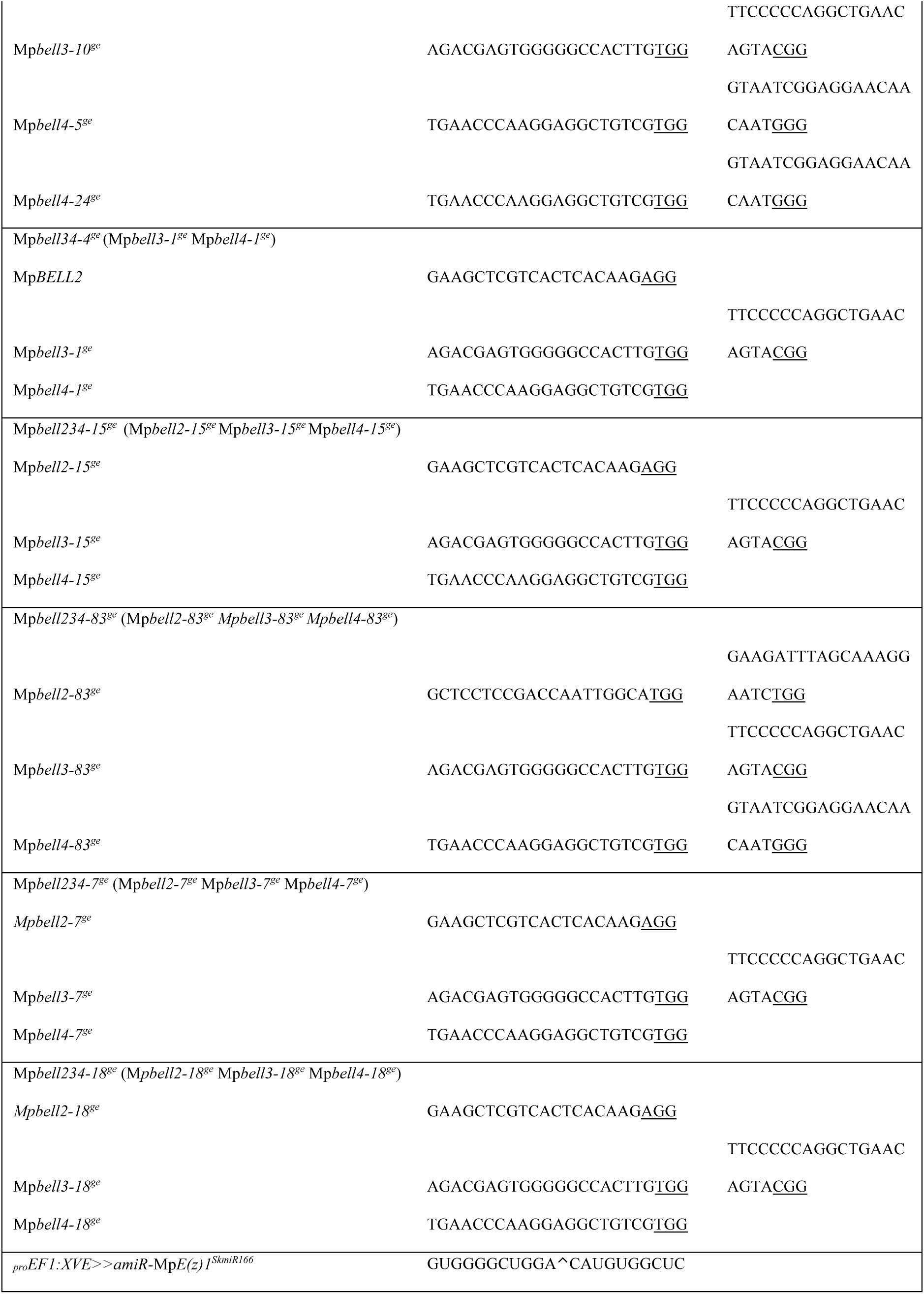
Guide RNA sequences used to generate mutant alleles.

### GUS staining

Generally, detection of GUS activity in a minimum of three independent lines was performed overnight at 37°C in GUS staining solution (0.05 M NaPO4, pH 7.2, 2 mM K3[Fe(CN)6], 2 mM K4[Fe(CN)6], 0.3 % (v/v) Triton^TM^ X-100) supplemented with 0.6 mg/ml b-D-glucopyranosiduronic acid (X-gluc) dissolved in N,N-dimethylformamide (DMF). Only detection of *_pro_*Mp*KNOX1:GUS* activity in archegoniophores was performed for 4 h in staining solution containing 5 mM K3[Fe(CN)6] and 5 mM K4[Fe(CN)6]. Antheridiophores were cleared in 70% (w/v) chloral hydrate in 10% (v/v) glycerol solution.

### Phylogenetics

Predicted TALE HD sequences were assembled from land plants, charophytes and Chlorophytes, via Genbank or additional sources as detailed below. Gymnosperm sequences were obtained from Congenie (congenie.org), fern sequences from *Equisetum* and *Azolla* transcriptomes (de Vries et al., 2016, Vanneste et al., 2015), bryophyte sequences, other than *M. polymorpha* (Phytozome) and *P. patens* (Genbank), from available transcriptomes (Wickett et al., 2014), *Klebsormidium* and *Caulerpa* sequences derived from genome sequences (Hori et al., 2014, Arimoto et al., 2019), and other charophyte and Chlorophyte sequences from Genbank and additional transcriptomes (Cooper and Delwiche, 2016, Ju et al., 2015).

Complete or partial coding nucleotide sequences were manually aligned as amino acid translations using Se-Al v2.0a11 for Macintosh (http://tree.bio.ed.ac.uk/software/seal/). We excluded ambiguously aligned sequence to produce an alignment of 228 nucleotides (76 amino acids) for 82 BELL sequences (see Figure 2—figure supplement 1). Alignments of KNOX genes included the homeodomain, MEINOX (KNOX) and ELK domains (Joo et al., 2018), comprising 606 nucleotides (202 amino acids) for 103 KNOX sequences. Alignments of nucleotides and amino acids were employed in subsequent Bayesian analysis. Bayesian phylogenetic analysis was performed using Mr. Bayes 3.2.1 (Huelsenbeck and Ronquist, 2001, Huelsenbeck et al., 2001). Analyses were performed on both the nucleotide and amino acid alignments. The fixed rate model option JTT + I was used based on analysis of the alignments with ProTest 2.4 (Abascal et al., 2005). The Bayesian analyses for the nucleotide data sets were run for 5,000,000 (BELL) or 1,000,000 (KNOX) generations, which was sufficient for convergence of the two simultaneous runs (BELL, 0.0418; KNOX, 0.0416). Analyses for the amino acid data sets were run for 4,000,000 (BELL) or 2,000,000 (KNOX) generations, which was sufficient for convergence of the two simultaneous runs (BELL, 0.037; KNOX, 0.039). In both cases, to allow for the burn-in phase, 50% of the total number of saved trees was discarded. The graphic representation of the trees was generated using the FigTree (version 1.4.0) software (http://tree.bio.ed.ac.uk/software/figtree/). Sequence alignments and command files used to run the Bayesian phylogenetic analyses provided upon request.

### RNA-seq

G2 gemmae were grown under normal growth conditions (23°C, 24h white light) on ½ B5 Gamborg’s media for 10 days, and then transferred via forceps to plates containing 5μM 17-β-estradiol. Plants were sampled at specific time points and immediately frozen in liquid nitrogen, then ground up using mortal and pestle. RNA was extracted from samples at time point 0 (not induced), plus 5 time points (3 h, 6 h, 9 h, 24 h, 48 h) from ≈100 mg wet weight as described above. Samples were stored at −80°C until shipped to BGI Tech Solutions (Hong Kong) on dry ice for sequencing. Mapping of FastQC reads to the Marchantia_polymorpha.mainGenome.fasta (3.1) was done by bowtie2 using the Galaxy platform (https://usegalaxy.org/) and Cufflinks assembled transcripts were based on the published transcriptome (Mpolymorphav3.1.primaryTrs.gff3). Relative transcript abundance was compared between un-induced transgenic lines and induced lines at different time points and visualized using IGV genome viewer (http://software.broadinstitute.org/software/igv/). We analysed the trajectory of de-repression of target genes and used the latest time point after induction (48h) since expression changes of identified target genes were highest at this time point.

### ChIP-seq

Wild-type and *_pro_EF1:XVE*>>*amiR-*Mp*E(z)^SkmiR166^* G2 were grown on ½ B5 media for 2 weeks. Induction of *_pro_EF1:XVE*>>*amiR-*Mp*E(z)^SkmiR166^* lines was done by flooding the plates with 50μM 17-β-estradiol and left for 48h before snap freezing in liquid nitrogen. Wild-type and un-induced plants were likewise snap frozen and all samples were shipped on dry ice to Active Motif, Inc. (USA; https://www.activemotif.com/catalog/820/histonepath) for H3K27me3 antibody qualification, ChIP-seq and sequencing of input DNA. Briefly, tissue was ground to a powder in liquid nitrogen and placed in 40 mL Plant Cross-linking and Extraction Buffer with addition of 1mL 37% formaldehyde and 400µL 0.1M PMSF and incubated, with stirring under a vacuum for 10 minutes. After addition of 1.6mL 2.5M Glycine, sample was mixed for an additional 5 minutes in a vacuum. Following centrifugation (10 minutes at 2500rpm, 4°C) the pellet was resuspended in ice-cold diH_2_O, and centrifugation was repeated with an ice-cold diH_2_O wash. Pellet was freeze dried for 30 minutes and stored at −80°C

Chromatin was prepped by resuspending freeze dried pellet in 5mL nuclei isolation buffer, 5µL protease inhibitor cocktail (PIC) and 50µL 0.1M PMSF. Samples were incubated on ice for 10 minutes and subsequently homogenized by 20 strokes of douncing. Following centrifugation (10 minutes at 2500rpm, 4°C) the pellet was resuspended in 500µL DOC sonication buffer, 5µL PIC and 5µL 0.1M PMSF, and incubated on ice for 10 minutes. Samples were sonicated for 10 minutes in 30 second pulses in the presence of glass beads. Following centrifugation, the supernatant was aliquoted and stored at −80°C.

Genomic DNA (Input) was prepared by treating aliquots of chromatin with RNase, proteinase K and heat for de-crosslinking, followed by ethanol precipitation. Pellets were resuspended and the resulting DNA was quantified on a NanoDrop spectrophotometer. Extrapolation to the original chromatin volume allowed quantitation of the total chromatin yield. An aliquot of chromatin (6 ug) was precleared with protein A agarose beads (Invitrogen). Genomic DNA regions of interest were isolated using 12 ug of antibody against H3K27me3 (Millipore, 07-449, Lot 2653203). Complexes were washed, eluted from the beads with SDS buffer, and subjected to RNase and proteinase K treatment. Crosslinks were reversed by incubation overnight at 65°C, and ChIP DNA was purified by phenol-chloroform extraction and ethanol precipitation.

Illumina sequencing libraries were prepared from the ChIP and Input DNAs by the standard consecutive enzymatic steps of end-polishing, dA-addition, and adaptor ligation. After a final PCR amplification step, the resulting DNA libraries were quantified and sequenced on Illumina’s NextSeq 500 (75 nt reads, single end). Reads were aligned to the *M. polymorpha* v3.1 genome using the BWA algorithm (v0.6.1-r104; default settings; (Li and Durbin, 2009)). Duplicate reads were removed and only uniquely mapped reads (mapping quality >= 25) were used for further analysis. Peak locations were determined using the MACS algorithm (v1.4.2; (Zhang et al., 2008)) with a cutoff of p-value = 1e-5, and wiggle files were generated by using the MACS –wig option (other MACS parameters were: mfold 8,30; --space 32; --nomodel; -- gsize=200000000).

### Data repository

The data discussed in this publication have been deposited in NCBI’s Gene Expression Omnibus (Edgar et al., 2002) and are accessible through GEO Series accession number GSE147756 (https://www.ncbi.nlm.nih.gov/geo/query/acc.cgi?acc=GSE147756).

---

**Figure 1—figure supplement 1.**
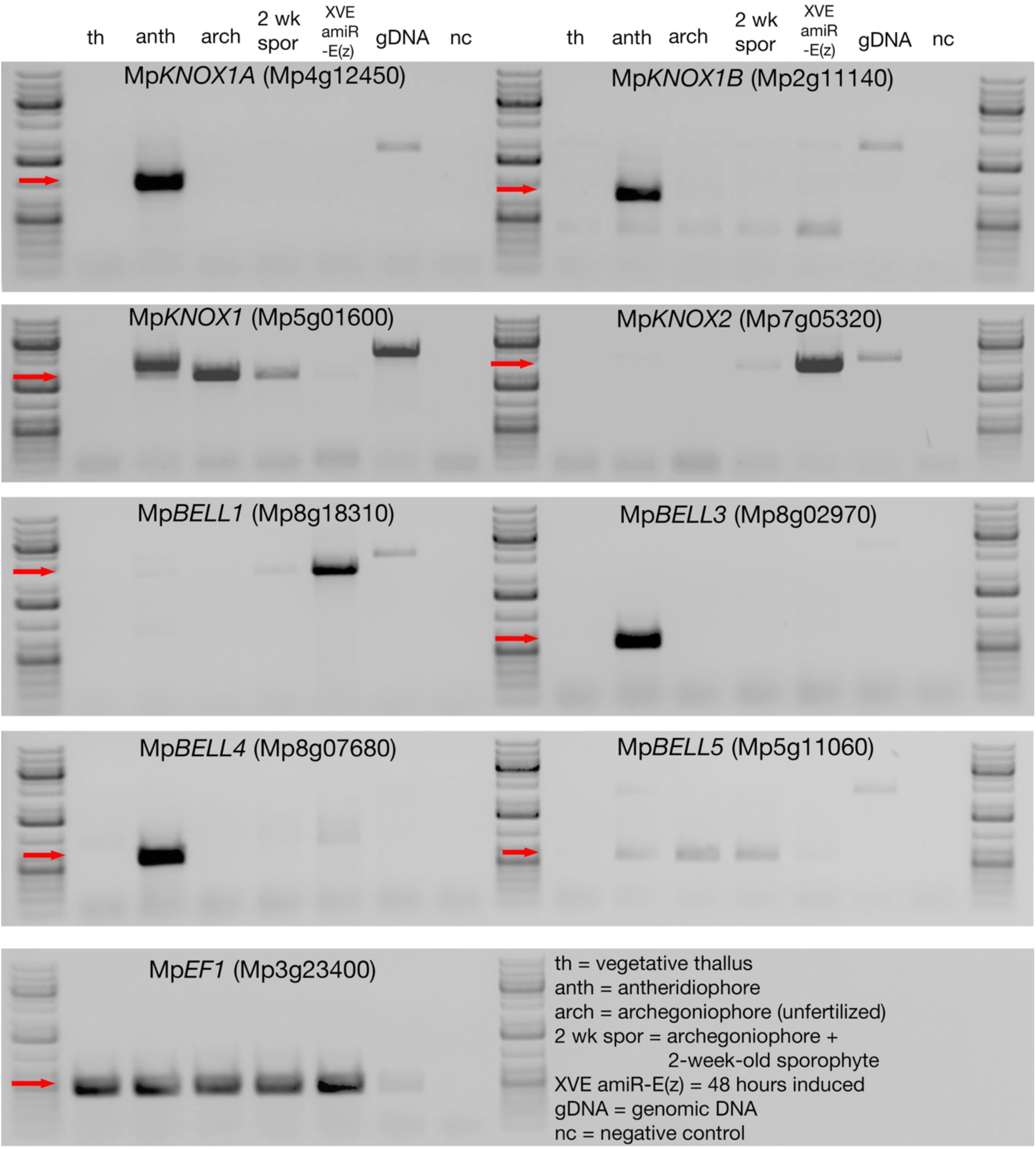
Expression of *M. polymorpha* TALE-HD genes. RNA extracted from various tissues of *M.polymorpha* ssp *ruderalis*, ecotype BoGa tissue were subjected to sqRT-PCR (30 cycles, except Mp*KNOX1* and Mp*KNOX2* at 32 cycles; 55°C anneal; 1 minute extension, except Mp*KNOX1*, Mp*KNOX2*, Mp*BELL1* at 2 minutes). The expected size band from cDNA is indicated by a red arrow at left of each panel. We were unable to amplify a product from Mp*BELL2*.

**Figure 1—figure supplement 2.**
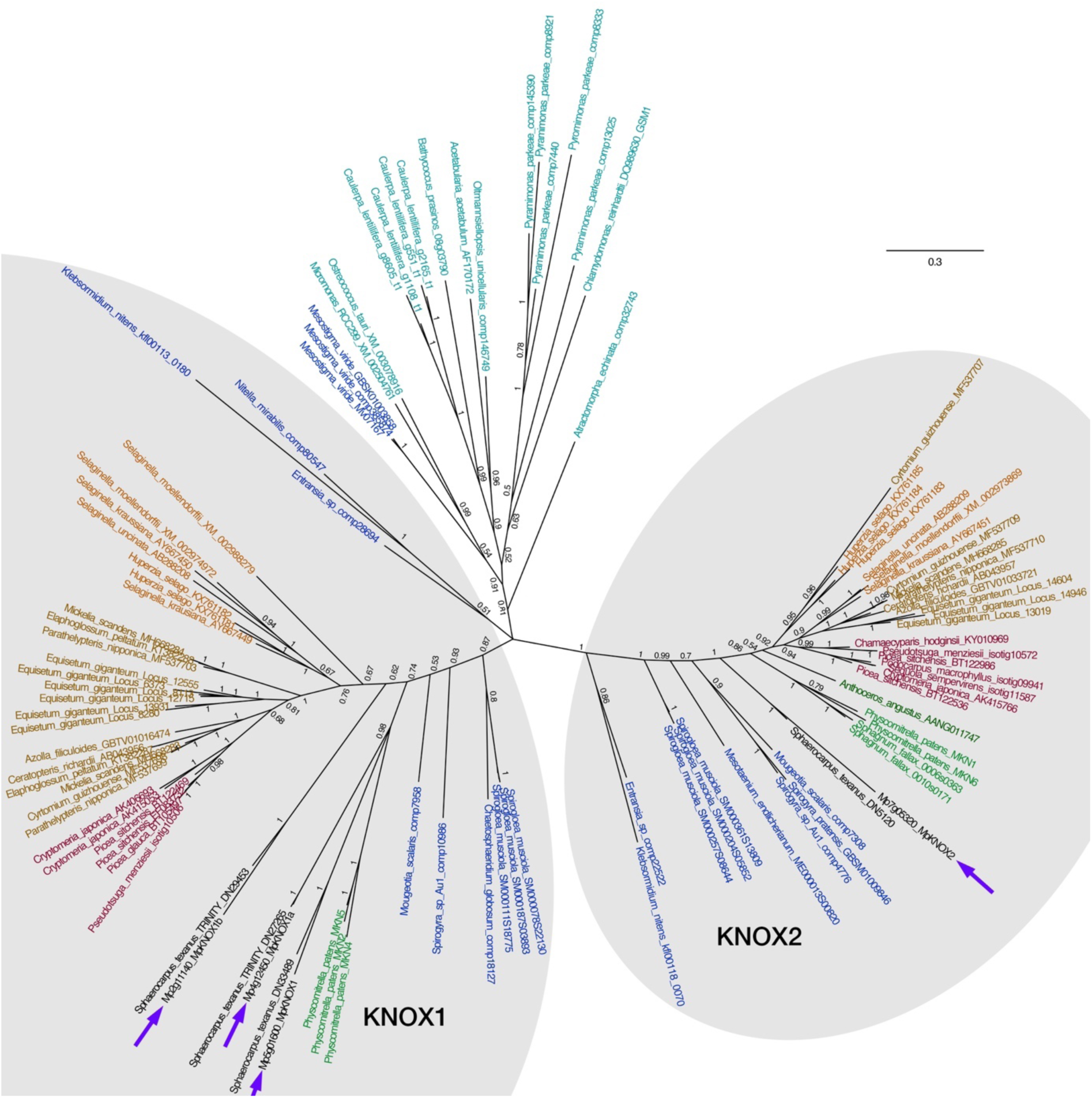
Unrooted Bayesian phylogram of Viridiplantae KNOX-TALE class homeodomain genes. Tree constructed using nucleotide alignment of the homeodomain and MEINOX and ELK domains; the phylogenetic tree constructed using amino acid characters was congruent. The four *M. polymorpha* KNOX genes are highlighted (purple arrows). The tree topology suggests a single KNOX gene in the common ancestor of the Viridiplantae, and possibly the Streptophyta, with a gene duplication in an ancestral charophycean alga after the divergence of *Mesostigma* producing ancestral KNOX1 and KNOX2 genes (Sakakibara, 2016, Joo et al., 2018, Frangedakis et al., 2017). The KNOX2 genes, including charophyte sequences, form a well-supported monophyletic clade (posterior probability = 1). The KNOX1 clade is less well-supported with regard to the early diverging charophyte sequences; however, land plant KNOX1 genes, including Mp*KNOX1*, share a conserved MEINOX domain (see also Figure 3—supplemental figure 1). The ancestral land plant likely had single KNOX1 and KNOX2 orthologs, a condition retained in *M. polymorpha*. Genes orthologous with Mp*KNOX1A* and Mp*KNOX1B* are found in *Sphaerocarpus texanus* indicating their origin at least as early as the basal Marchantiopsida. Chlorophyte, light blue; charophyte, dark blue; liverwort, black; moss, green; hornwort, dark green; lycophyte, brown, fern, orange; seed plant, red. Numbers indicate posterior probability values >50%.

**Figure 2—figure supplement 1.**
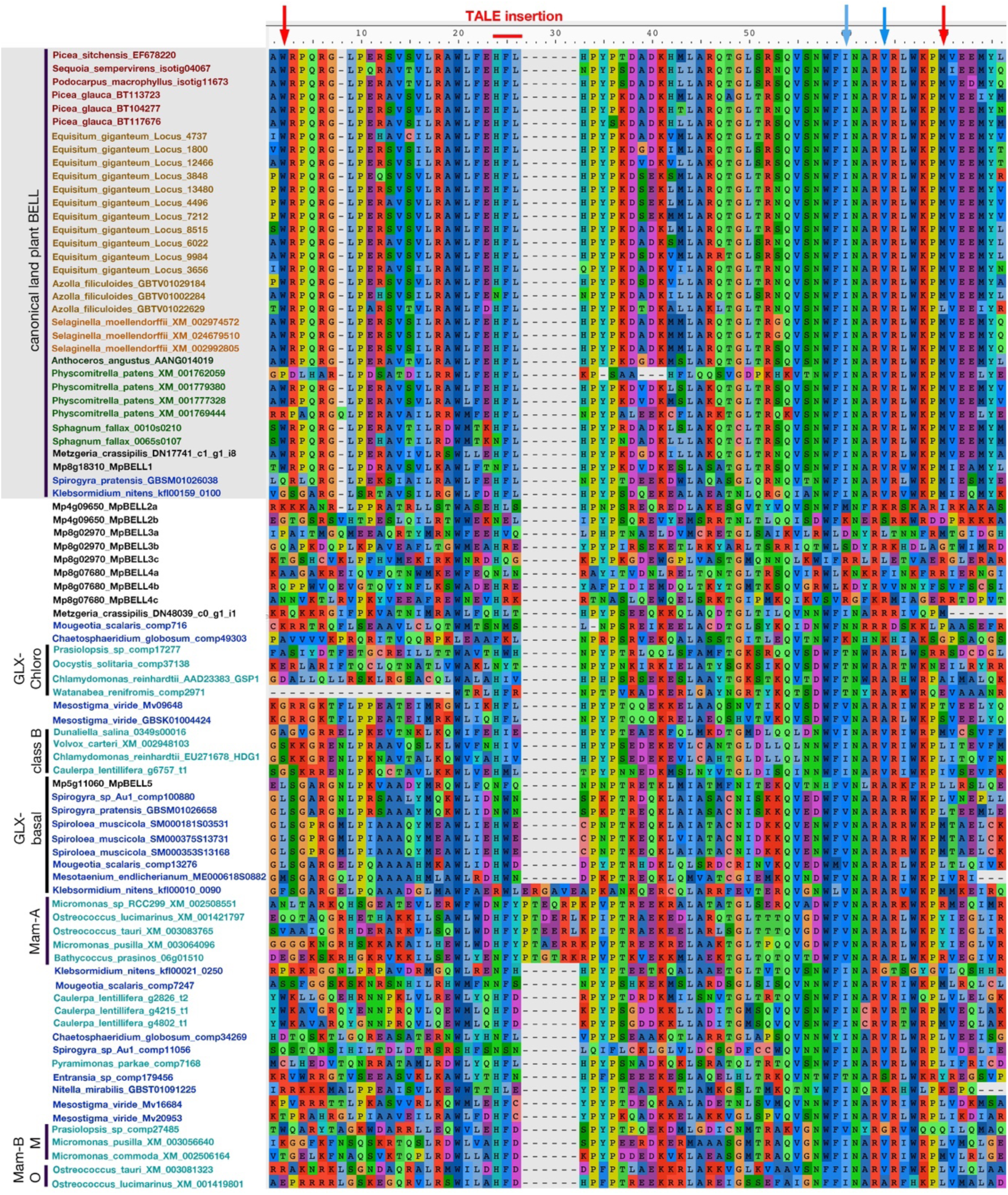
Alignment of Viridiplantae BELL homeodomains. This alignment was used in the construction of the phylogenetic tree in Figure 2. Homeodomain delimited by the red arrows. Canonical land plant BELL sequences are shaded light grey at the top; color coding of taxa as in Figure 2, with *M. polymorpha* and other liverwort sequences in black. Based on the identity of specific residues (at position 60 and 64 in the figure and highlighted by blue arrows) in the homeodomain Lee et al. (Lee et al., 2008) suggested that *C. reinhardtii HDG1* was orthologous to canonical land plant BELL genes (characterized by I and V residues at the positions); this is supported by phylogenetic analysis (Figure 2). The second (b) and third (c) homeodomains of Mp*BELL2*, Mp*BELL3*, and Mp*BELL4* are progressively more divergent that the carboxyl-most (a) homeodomain

**Figure 4—figure supplement 1.**
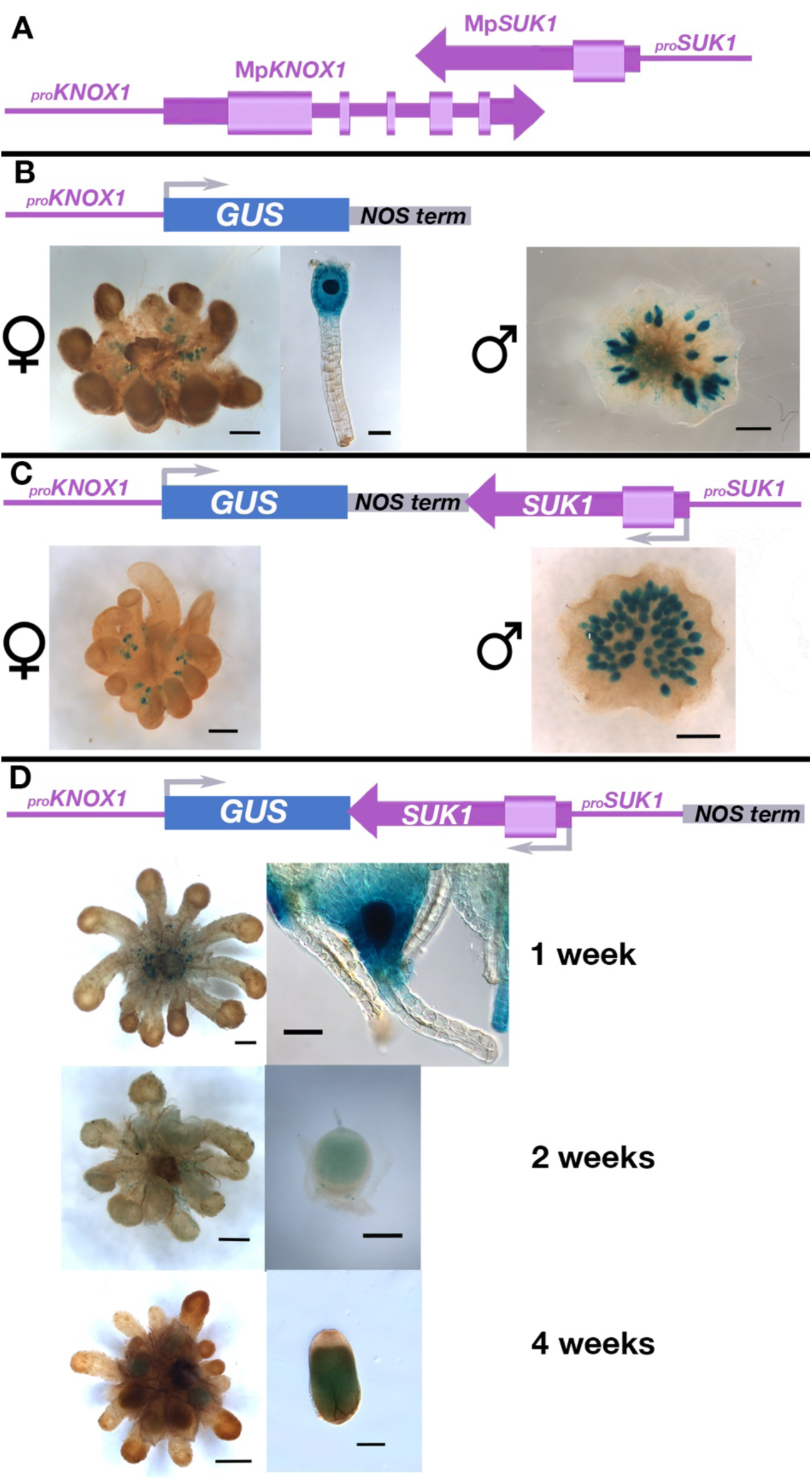
Expression profile of *_pro_*Mp*KNOX1* transcriptional β-glucuronidase reporter gene fusions unravel regulation by antisense transcript Mp*SUK1.* Sequences in purple are from the endogenous locus, while sequences in other colors are heterologous. (A) Endogenous Mp*KNOX1*/Mp*SUK1* locus. (**B**) A construct including 4.6 kb sequence upstream (5’) of the Mp*KNOX1* translational start site fused with the GUS reporter gene followed by a NOS transcriptional terminator (250 bp) drives expression in both the egg cell (left) and antheridia (right). (**C**) Inclusion of the Mp*SUK1* locus, including 1.5 kb upstream (5’) of the Mp*SUK1* translational start site, downstream of the NOS terminator results in the same expression pattern, i.e. in both the egg cell and antheridia. **(D)** Only in constructs featuring an assembly in which *_pro_*Mp*KNOX1:GUS* and *_pro_*Mp*SUK1:*Mp*SUK1* are opposing and not interrupted by a transcriptional terminator is expression of Mp*KNOX1* restricted to the female egg cell (Figure 4N-P) and developing sporophyte, shown here at 1 week, 2 weeks, and 4 weeks post fertilization. Bar = 50 µm (B, G) 250 µm (I, K) 500 µm (A, D) 1000 µm (C, E, F, H, J).

**Figure 4—figure supplement 2.**
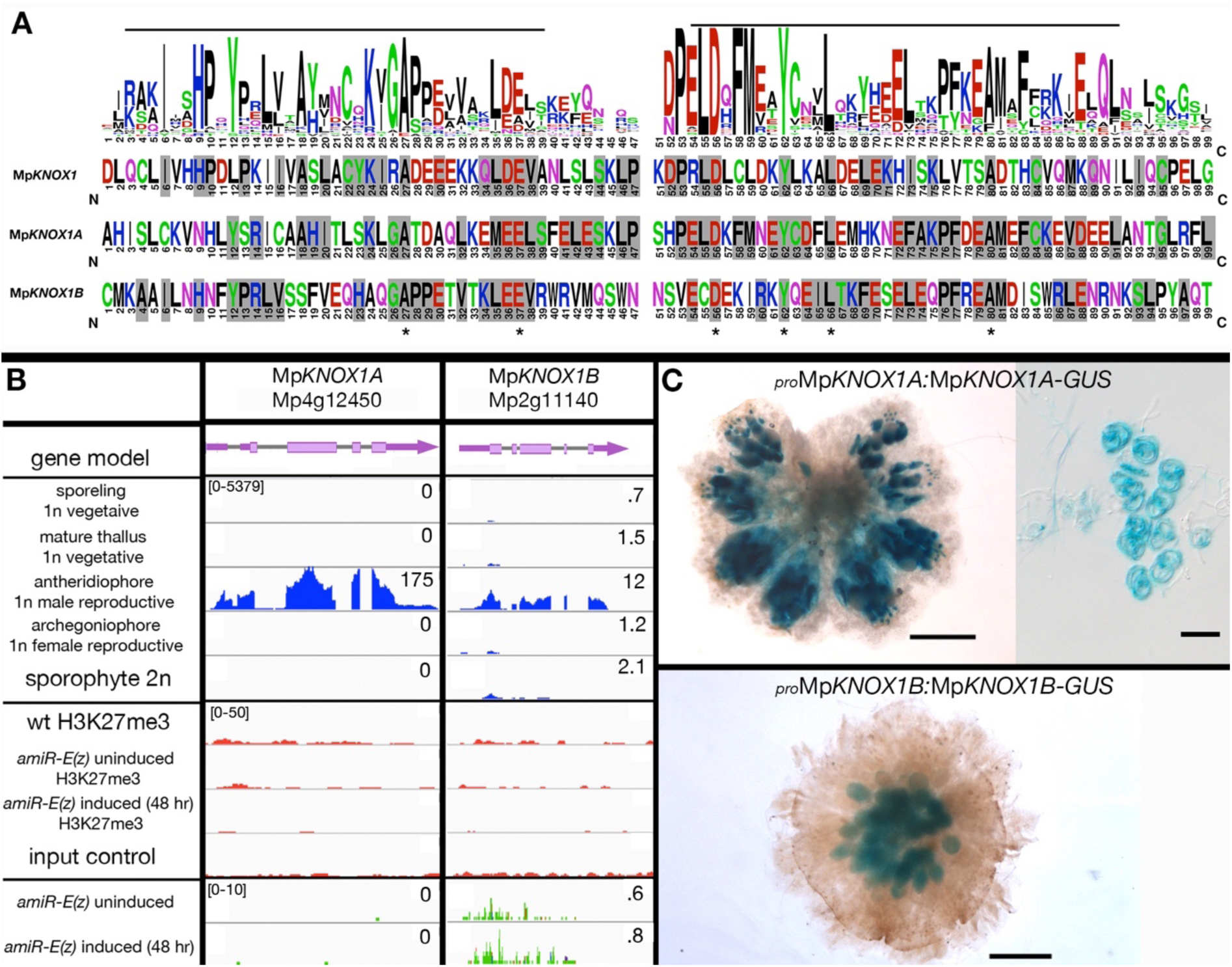
Mp*KNOX1A* and Mp*KNOX1B* lack the homeodomain and are both primarily expressed in the antheridiophores. (**A**) **Alignment of MpKNOX1 MEINOX domains with land plant KNOX1 proteins.** A consensus sequence of the MEINOX domain (Burglin, 1997, Crooks et al., 2004) of land plant KNOX1 proteins was computed from sequences in Figure 1—figure supplement 1, with the alignment of the three *M. polymorpha* KNOX1 proteins displayed below. The lines above the sequence highlight the previously defined KNOX subdomains of the MEINOX domain. Grey shading indicates amino acids conserved with the land plant consensus and asterisks highlight amino acids conserved in all three *M. polymorpha* KNOX1 sequences. (**B**) Panels as described in Figure 3. Expression of both genes is predominantly detected in the antheridiophore, with neither gene being marked with H3K27me3 in the vegetative gametophyte and nor up-regulated in the *amiR-*Mp*E(z)1* background. (**C**) Translational fusion reporter line expression containing 1kb (Mp*KNOX1A*) and 2.3kb (Mp*KNOX1B*) of the putative 5’ regulatory sequences suggests that the antheridiophore expression identified by RNAseq is primarily in the antheridia. Expression of a translational fusion of MpKNOX1A with GUS was detectable in the sperm. Weak Mp*KNOX1A* reporter line expression was also found in the tissues of the archegoniophore and archegonium. Bars = 200µm (antheridiophores) and 10µm (sperm).

**Figure 5—figure supplement 1.**
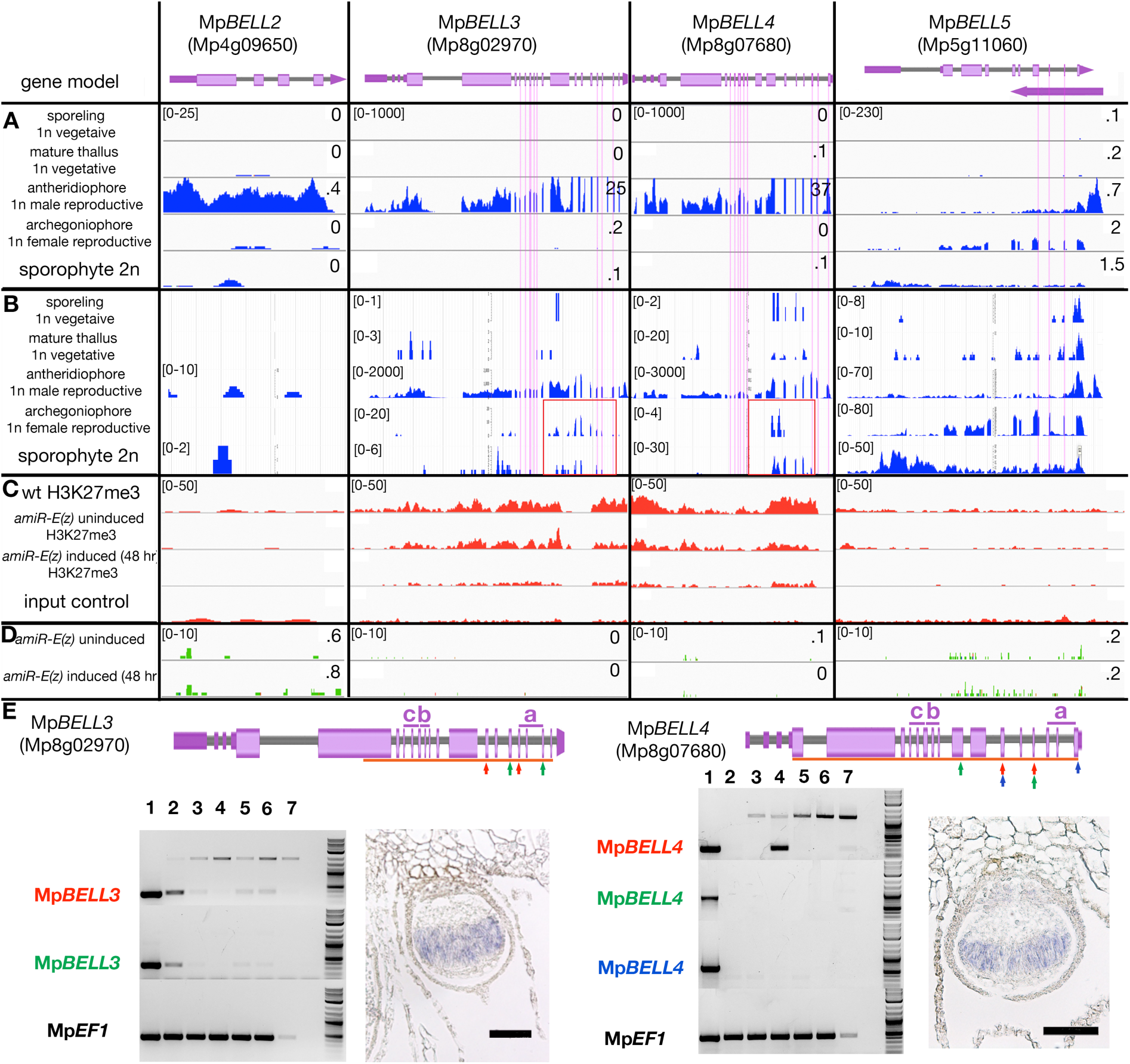
Expression profiles and H3K27me3 marks at MpBELL loci. Panels as described in Figure 3. Vertical pink lines highlight exons encoding the homeodomains, each spanning three exons, in Mp*BELL3*, Mp*BELL4* and Mp*BELL5*. (**A**) Expression in different tissues of wild-type *M. polymorpha* based on RNA-seq experiments (Bowman et al., 2017). (**B**) Strand-specific expression at each of the MpBELL loci (marchantia.info). Potential alterative shorter transcripts (highlighted in red boxes) are detected for Mp*BELL3* in archegoniophores and at a lower level, in sporophytes, and for Mp*BELL4* in sporophytes. (**C**) Both Mp*BELL3* and Mp*BELL4* are marked by H3K27me3 in vegetative gametophytes. Mp*BELL2* and Mp*BELL5* are not marked by H3K27me3 in vegetative gametophytes. (**D**) None of the MpBELL genes, whether marked by H3K27me3 or not, are not induced by loss of Mp*E(z)* activity. (**E**) Positions of homeodomains are delineated above gene model. rtPCR experiments using color-coded primer pairs. 1, wild-type antheridiophores; 2, wild-type archegoniophores (not fertilized); 3, wild-type 1 week post-fertilization (wpf); 4, wild-type 3 wpf; 5, wild-type female x Mp*bell34-4* (1wpf); 6. wild-type female x Mp*bell2/3/4-15* (1wpf); 7, genomic DNA; 8. no template control; size marker on far right. *In situ* probes spanned exons indicated by orange bars. Shown are sporophytes 3 weeks post-fertilization; bars = 100µm. All scales (FPKM), except those in the strand-specific expression panels, are set to a standard within each outlined box; all H3K27me3 panels set to 50; for RNA-seq data the FPKM values (if less than 10, rounded to .1) are presented. Scales in the strand-specific expression panels vary according to tissue to allow potential weakly-expressed transcripts to be visualized.

**Figure 6—figure supplement 1.**
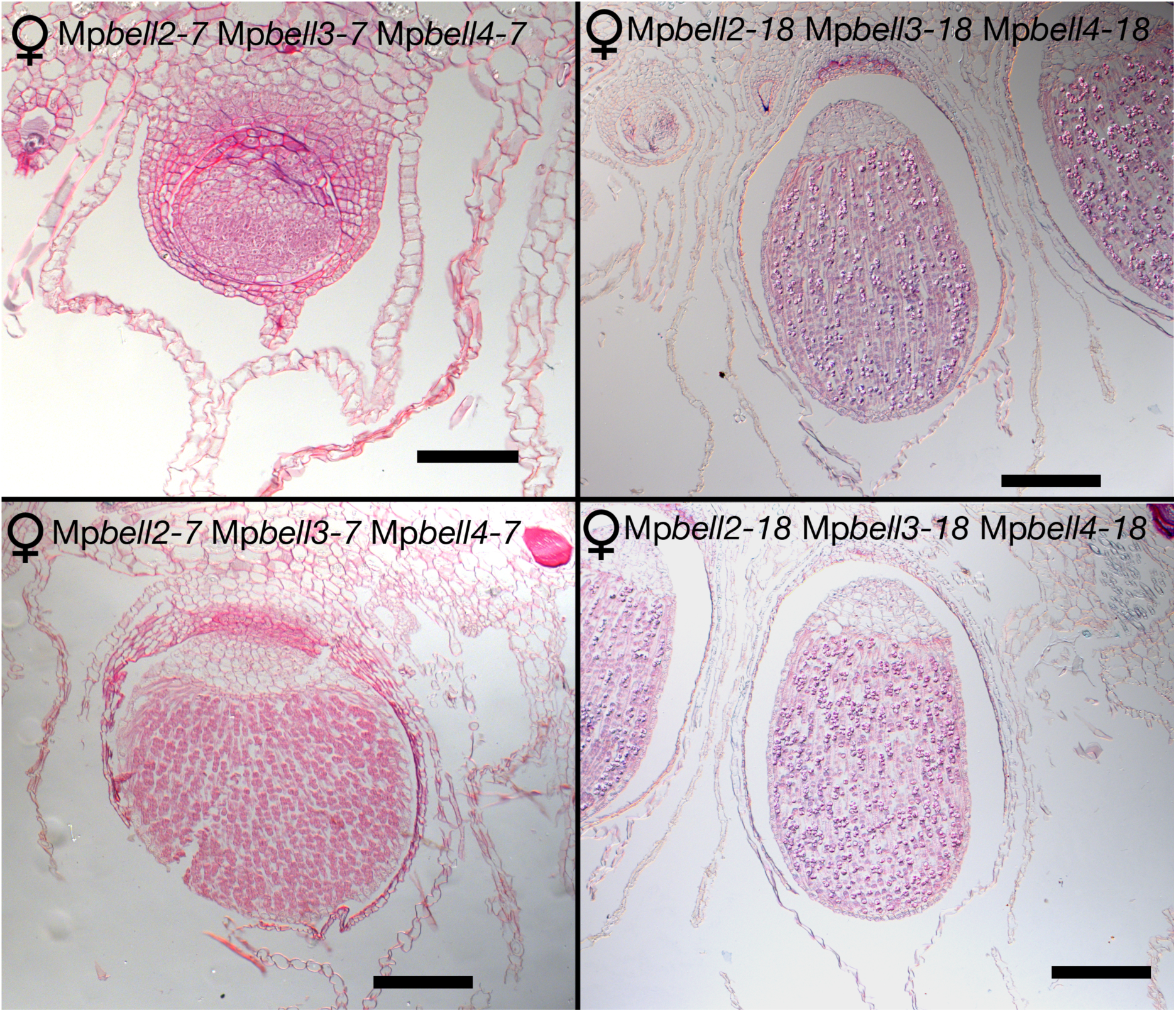
Mature sporophytes are produced in Mp*bell234* females. Sporophytes develop normally following crosses between a female Mp*bell234-7* triple mutant or a female Mp*bell234-18* triple mutant and wild-type male plants. Sporophytes depicted were observed 2-3 weeks (Mp*bell234-7*) or 3-4 weeks (Mp*bell234-18*) post fertilization. Bar = 100 µm.

**Figure 6—figure supplement 2.**
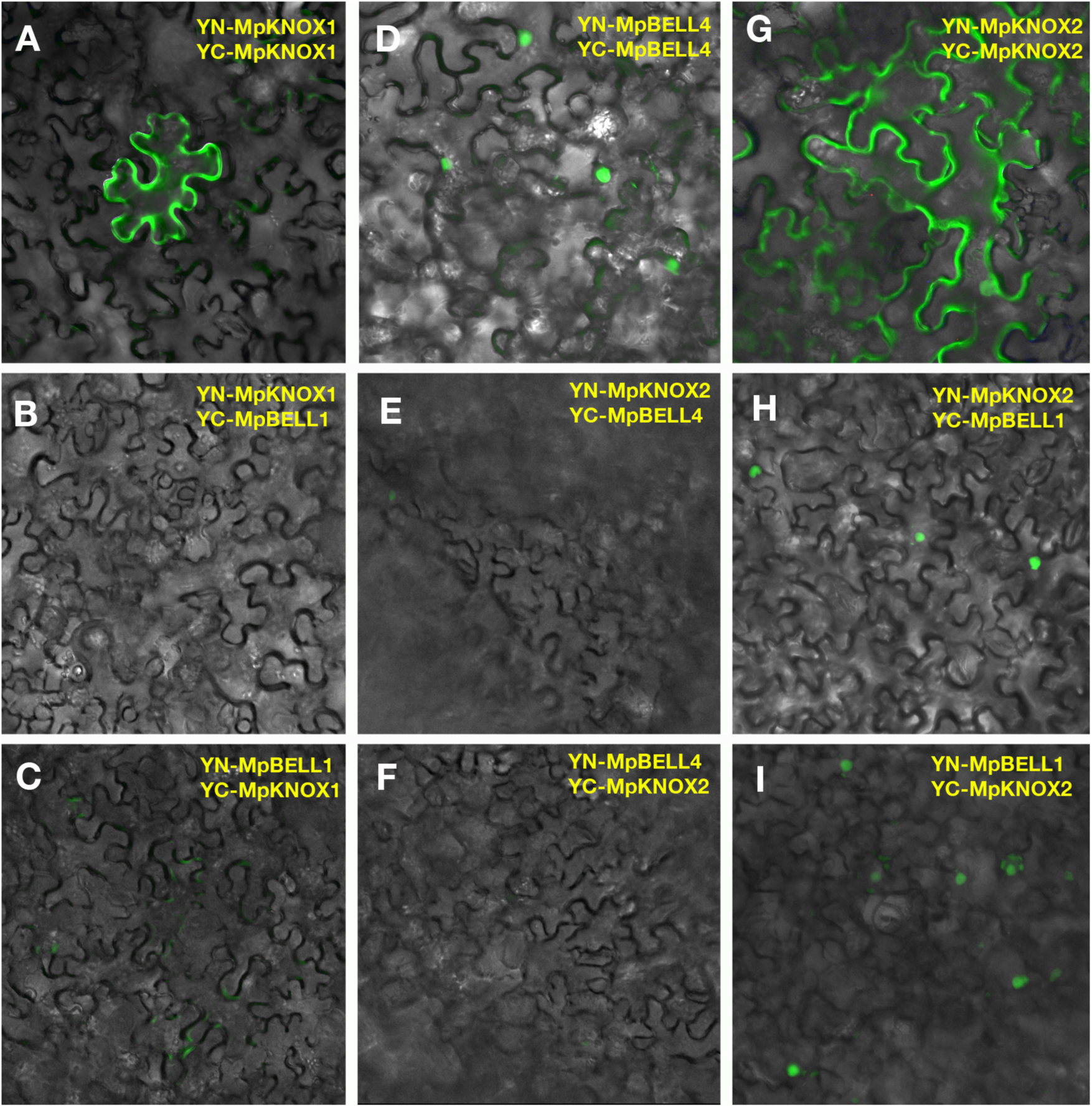
MpKNOX1/MpBELL4 and MpKNOX2/MpBELL1 exhibit specificity in their interactions. BiFC assay of protein-protein interaction in *N. benthamiana* leaves. (A) MpKNOX1 homodimers show cytoplasmic localisation. (B-C) No interaction can be observed when MpKNOX1 is co-expressed with MpBELL1. (D) MpBELL4 homodimers localizes to the nucleus. (E-F) Co-expression of both MpBELL4 and MpKNOX2 show no interaction. (G) MpKNOX2 homodimers alone show cytoplasmic localisation. MpBELL1 homodimers locate to the nucleus (data not shown). (H-I) However, when MpKNOX2 and MpBELL1 are co-expressed, interaction signal is located to the nucleus. Note that (A) and (B) are taken from Figure 6B and C for reference.

**Figure 6—figure supplement 3.**
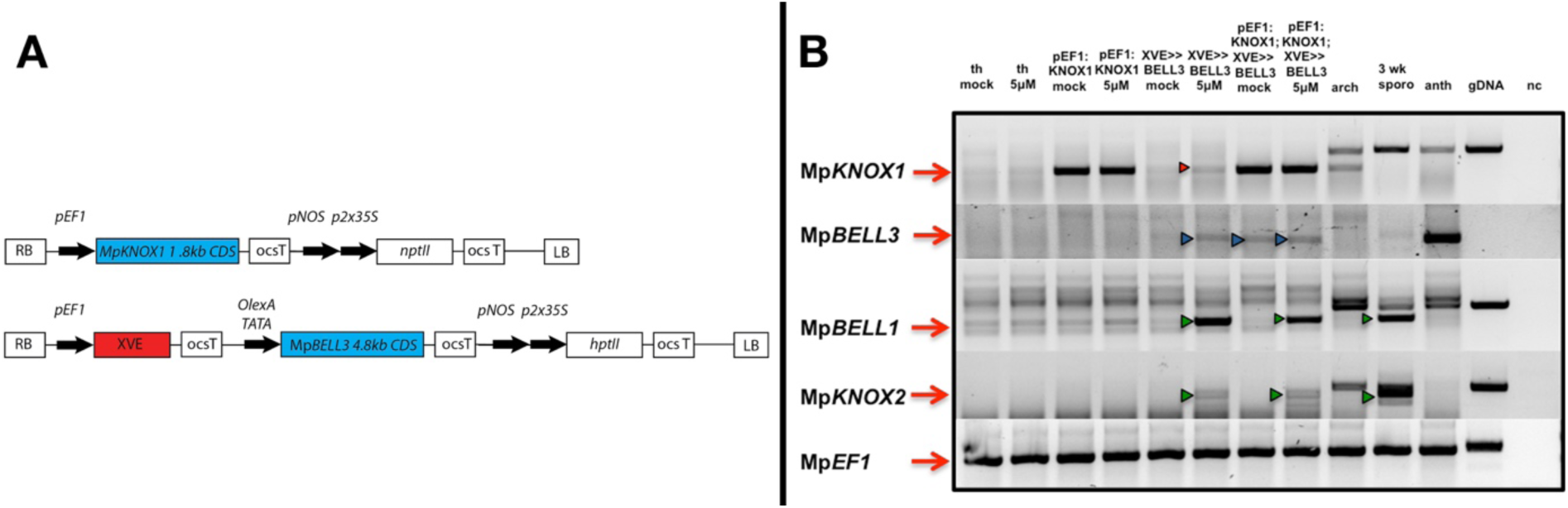
Co-expression strategy to express Mp*KNOX1* and Mp*BELL3* simultaneously in the vegetative gametophyte. (A) Plants were doubly transformed using one construct *in cis* and one *in trans*: *_pro_EF1:*Mp*KNOX1* constitutively expressing Mp*KNOX1* CDS and *_pro_EF1:XVE>>*Mp*BELL3* expressing a 4.8kb fragment of the putative 9kb CDS after induction. Note that the pair of constructs features two different selection markers (*nptII* for selection on G418 and *hptII* for hygromycin selection) for selection of doubly transformants. (B) sqRT-PCR of mock and 5 μM 17-β-estradiol treated plants after 72h induction. Mp*KNOX1* transcription is induced after 72h in *_pro_EF1:XVE>>*Mp*BELL3* lines (red arrowhead). Antheridia specific Mp*BELL3* expression is induced in *_pro_EF1:XVE>>*Mp*BELL3* lines, in case of the doubly transformants also in the un-induced line (blue arrowheads). Mp*KNOX2* and Mp*BELL1*, both normally expressed primarily in the sporophyte are de-repressed after induction via Mp*BELL3* expression (green arrowheads). th = thallus; arch = archegoniophore; 3wk sporo = archegoniophore including sporophytes 3 weeks past fertilization; anth = antheridiophores; gDNA = genomic DNA; nc = negative control.

**Figure 7—figure supplement 1.**
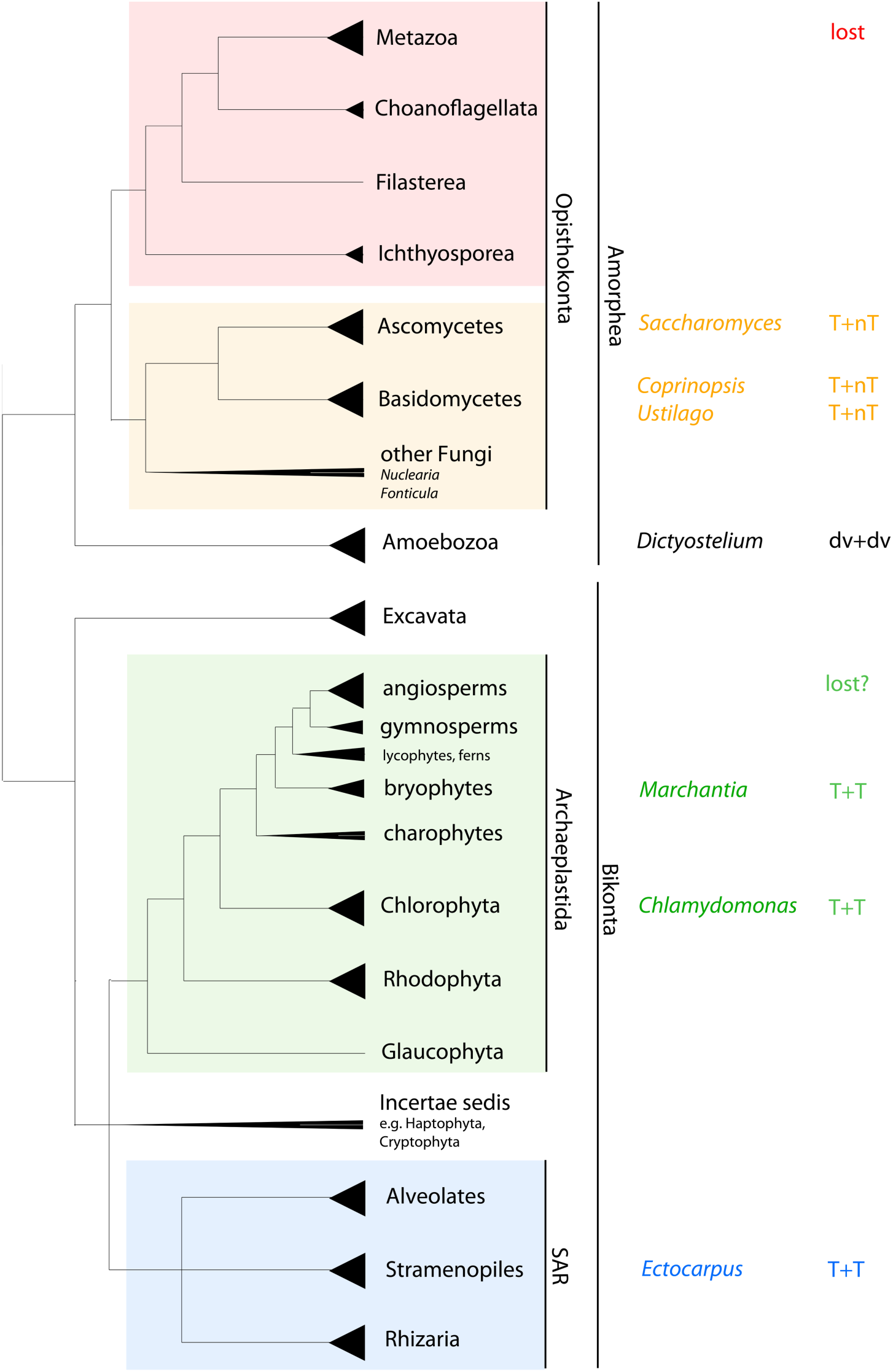
Phylogenetic perspective on HD gene mediated haploid to diploid transition in eukaryotes. Unrooted phylogenetic tree adapted from Adl et al. (Adl et al., 2012) and Keeling et al. (Keeling et al., 2014). Triangles at end of branches represent monophyletic groups; triangles extending to the node represent paraphyletic groups. Taxa in which functional data are available are listed, as is the mode of HD involvement. While the ancestral eukaryote possessed both TALE and non-TALE HD genes, the predicted composition of the heterodimer in the ancestral eukaryote is equivocal. T = TALE-HD; nt = non-TALE-HD; dv = divergent HD.

**Table 1—figure supplement 1.**
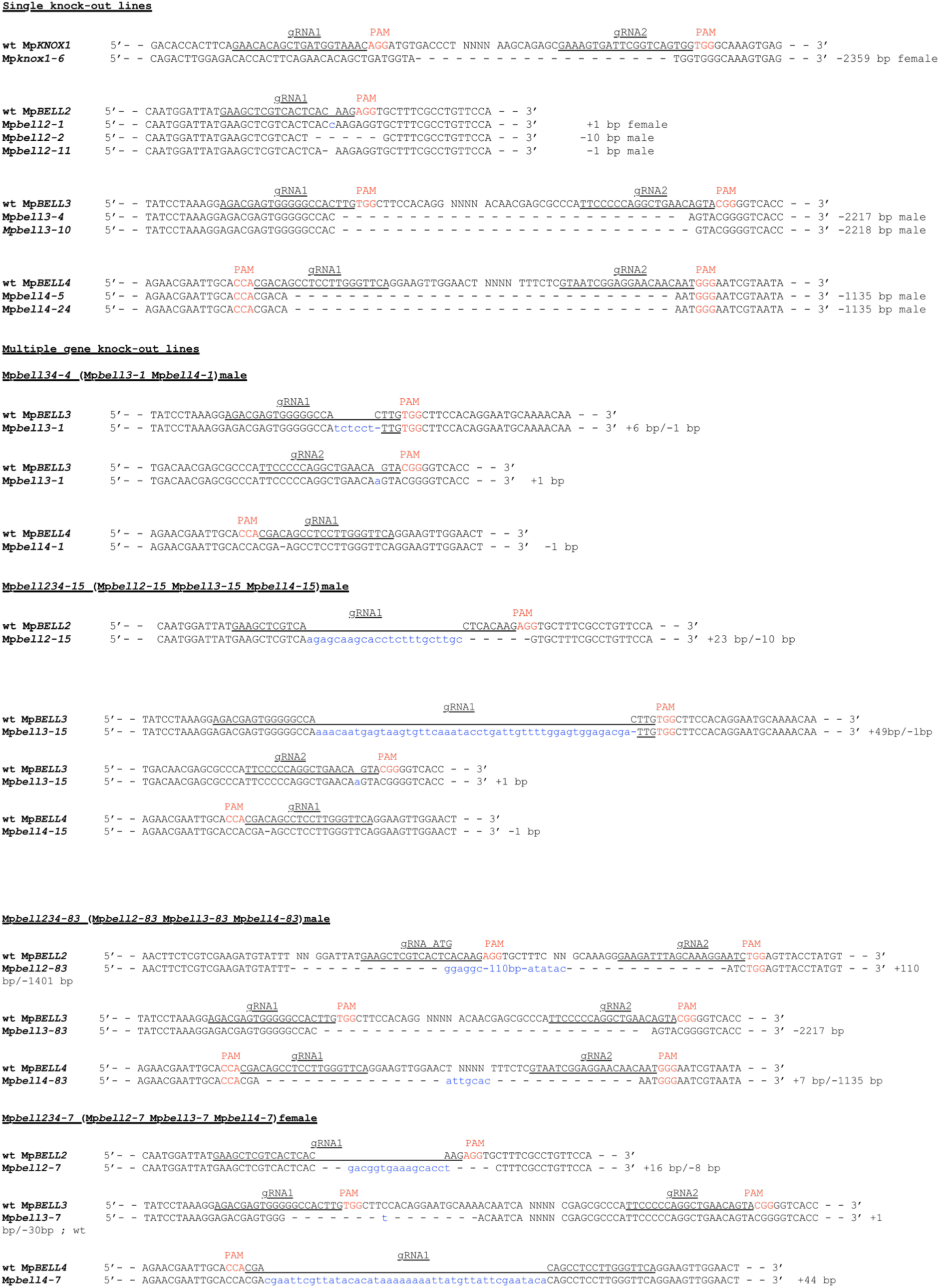
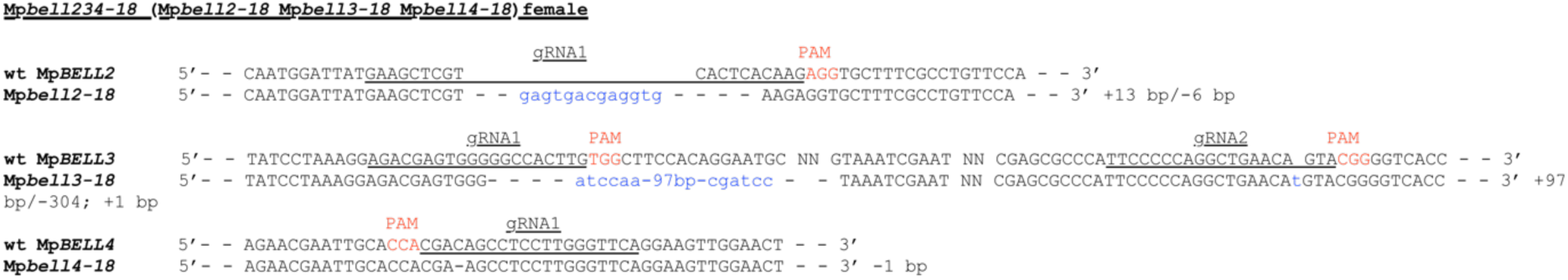
Molecular lesions of alleles listed in Table 1.

## Literature cited

Abascal, F., Zardoya, R. & Posada, D. 2005. ProtTest: selection of best-fit models of protein evolution.. Bioinformatics, 21, 2104–2105.

Adl, S. M., Simpson, A. G. B., Lane, C. E., Lukes, J., Bass, D., Bowser, S. S., Brown, M. W., Burki, F., Dunthorn, M., Hampl, V., Heiss, A., Hoppenrath, M., Lara, E., Le Gall, L., Lynn, D. H., Mcmanus, H., Mitchell, E. A. D., Mozley-Stanridge, S. E., Parfrey, L. W., Pawlowski, J., Rueckert, S., Shadwick, L., Schoch, C. L., Smirnov, A. & Spiegel, F. W. 2012. The Revised Classification of Eukaryotes. Journal of Eukaryotic Microbiology, 59, 429–493.

Althoff, F., Kopischke, S., Zobell, O., Ide, K., Ishizaki, K., Kohchi, T. & Zachgo, S. 2014. Comparison of the Mpef1α and Camv35 promoters for application in Marchantia polymorpha overexpression studies. Transgenic Research, 23, 234–244.

Arimoto, A., Nishitsuji, K., Higa, Y., Arakaki, N., Hisata, K., Shinzato, C., Satoh, N. & Shoguchi, E. 2019. A siphonous macroalgal genome suggests convergent functions of homeobox genes in algae and land plants. Dna Research, 26, 183–192.

Arun, A., Coelho, S. M., Peters, A. F., Bourdareau, S., Péres, L., Scornet, D., Strittmatter, M., Lipinska, A. P., Yao, H., Godfroy, O., Montecinos, G. J., Avia, K., Macaisne, N., Troadec, C., Bendahmane, A. & Cock, J. M. 2019. Convergent recruitment of Tale homeodomain life cycle regulators to direct sporophyte development in land plants and brown algae. eLife, 8, e43101.

Bellaoui, M., Pidkowich, M. S., Samach, A., Kushalappa, K., Kohalmi, S. E., Modrusan, Z., Crosby, W. L. & Haughn, G. W. 2001. The Arabidopsis Bell1 and Knox Tale homeodomain proteins interact through a domain conserved between plants and animals. Plant Cell, 13, 2455–2470.

Bharathan, G., Janssen, B. J., Kellogg, E. A. & Sinha, N. 1997. Did homeodomain proteins duplicate before the origin of angiosperms, fungi, and metazoa? Pnas, 94, 13749–13753.

Bower, F. O. 1908. Origin of a land flora: a theory based on the facts of alternation, London, MacMillan.

Bowman, J. L. 2016. A Brief History of Marchantia from Greece to Genomics. Plant and Cell Physiology, 57, 210–229.

Bowman, J. L., Araki, T., Arteaga-Vazquez, M. A., Berger, F., Dolan, L., Haseloff, J., Ishizaki, K., Kyozuka, J., Lin, S. S., Nagasaki, H., Nakagami, H., Nakajima, K., Nakamura, Y., Ohashi-Ito, K., Sawa, S., Shimamura, M., Solano, R., Tsukaya, H., Ueda, T., Watanabe, Y., Yamato, K. T., Zachgo, S. & Kohchi, T. 2016a. The Naming of Names: Guidelines for Gene Nomenclature in Marchantia. Plant and Cell Physiology, 57, 257–261.

Bowman, J. L., Kohchi, T., Yamato, K. T., Jenkins, J., Shu, S., Ishizaki, K., Yamaoka, S., Nishihama, R., Nakamura, Y., Berger, F., Adam, C., Aki, S. S., Althoff, F., Araki, T., Arteaga-Vazquez, M. A., Balasubrmanian, S., Barry, K., Bauer, D., Boehm, C. R., Briginshaw, L., Caballero-Perez, J., Catarino, B., Chen, F., Chiyoda, S., Chovatia, M., Davies, K. M., Delmans, M., Demura, T., Dierschke, T., Dolan, L., Dorantes-Acosta, A. E., Eklund, D. M., Florent, S. N., Flores-Sandoval, E., Fujiyama, A., Fukuzawa, H., Galik, B., Grimanelli, D., Grimwood, J., Grossniklaus, U., Hamada, T., Haseloff, J., Hetherington, A. J., Higo, A., Hirakawa, Y., Hundley, H. N., Ikeda, Y., Inoue, K., Inoue, S.-I., Ishida, S., Jia, Q., Kakita, M., Kanazawa, T., Kawai, Y., Kawashima, T., Kennedy, M., Kinose, K., Kinoshita, T., Kohara, Y., Koide, E., Komatsu, K., Kopischke, S., Kubo, M., Kyozuka, J., Lagercrantz, U., Lin, S.-S., Lindquist, E., Lipzen, A. M., Lu, C.-W., Luna, E. D., Martienssen, R. A., Minamino, N., Mizutani, M., Mizutani, M., Mochizuki, N., Monte, I., Mosher, R., Nagasaki, H., Nakagami, H., Naramoto, S., Nishitani, K., Ohtani, M., Okamoto, T., Okumura, M., Phillips, J., Pollak, B., Reinders, A., Rövekamp, M., Sano, R., Sawa, S., Schmid, M. W., Shirakawa, M., Solano, R., Spunde, A., Suetsugu, N., Sugano, S., Sugiyama, A., Sun, R., Suzuki, Y., Takenaka, M., et al. 2017. Insights into Land Plant Evolution Garnered from the Marchantia polymorpha Genome. Cell, 171, 287–304.

Bowman, J. L., Sakakibara, K., Furumizu, C. & Dierschke, T. 2016b. Evolution in the cycles of life. Annual Review of Genetics, 50, 133–154.

Bracha-Drori, K., Shichrur, K., Katz, A., Oliva, M., Angelovici, R., Yalovsky, S. & Ohad, N. 2004. Detection of protein-protein interactions in plants using bimolecular fluorescence complementation. Plant Journal, 40, 419–427.

Burglin, T. R. 1997. Analysis of Tale superclass homeobox genes (Meis, Pbc, Knox, Iroquois, Tgif) reveals a novel domain conserved between plants and animals. Nucleic Acids Research, 25, 4173–4180.

Cao, R., Wang, L. J., Wang, H. B., Xia, L., Erdjument-Bromage, H., Tempst, P., Jones, R. S. & Zhang, Y. 2002. Role of histone H3 lysine 27 methylation in polycomb-group silencing. Science, 298, 1039–1043.

Conner, J. A., Mookkan, M., Huo, H., Chae, K. & Ozias-Akins, P. 2015. A parthenogenesis gene of apomict origin elicits embryo formation from unfertilized eggs in a sexual plant. Proceedings of the National Academy of Sciences of the United States of America, 112, 11205–11210.

Conner, J. A., Podio, M. & Ozias-Akins, P. 2017. Haploid embryo production in rice and maize induced by Psasgr-Bbml transgenes. Plant Reproduction, 30, 41–52.

Cooper, E. & Delwiche, C. 2016. Green algal transcriptomes for phylogenetics and comparative genomics. figshare.https://dx.doi.org/10.6084/m9.figshare.1604778.

Crooks, G. E., Hon, G., Chandonia, J. M. & Brenner, S. E. 2004. WebLogo: A sequence logo generator. Genome Research, 14, 1188–1190.

Czermin, B., Melfi, R., Mccabe, D., Seitz, V., Imhof, A. & Pirrotta, V. 2002. Drosophila enhancer of Zeste/Esc complexes have a histone H3 methyltransferase activity that marks chromosomal polycomb sites. Cell, 111, 185–196.

De Vries, J., Fischer, A. M., Roettger, M., Rommel, S., Schluepmann, H., Brautigam, A., Carlsbecker, A. & Gould, S. B. 2016. Cytokinin-induced promotion of root meristem size in the fern Azolla supports a shoot-like origin of euphyllophyte roots. New Phytologist, 209, 705–720.

Derelle, R., Lopez, P., Le Guyader, H. & Manuel, M. 2007. Homeodomain proteins belong to the ancestral molecular toolkit of eukaryotes. Evolution & Development, 9, 212–219.

Di Croce, L. & Helin, K. 2013. Transcriptional regulation by Polycomb group proteins. Nature Structural & Molecular Biology, 20, 1147–1155.

Edgar, R., Domrachev, M. & Lash, A. E. 2002. Gene Expression Omnibus: Ncbi gene expression and hybridization array data repository. Nucleic Acids Research, 30, 207–210.

Edwards, K., Johnstone, C. & Thompson, C. 1991. A Simple and Rapid Method for the Preparation of Plant Genomic Dna for Pcr Analysis. Nucleic Acids Research, 19, 1349–1349.

Ferris, P. J. & Goodenough, U. W. 1987. Transcription of novel genes, including a gene linked to the mating-type locus, induced by Chlamydomonas fertilization. Molecular Cellular Biology, 7, 2360–2366.

Fischle, W., Wang, Y. M. & Allis, C. D. 2003. Histone and chromatin cross-talk. Current Opinion in Cell Biology, 15, 172–183.

Flores-Sandoval, E., Dierschke, T., Fisher, T. J. & Bowman, J. L. 2016. Efficient and Inducible Use of Artificial Micrornas in Marchantia polymorpha. Plant and Cell Physiology, 57, 281–290.

Flores-Sandoval, E., Eklund, D. M. & Bowman, J. L. 2015. A Simple Auxin Transcriptional Response System Regulates Multiple Morphogenetic Processes in the Liverwort Marchantia polymorpha. Plos Genetics, 11, e1005207.

Frangedakis, E., Saint-Marcoux, D., Moody, L. A., Rabbinowitsch, E. & Langdale, J. A. 2017. Nonreciprocal complementation of Knox gene function in land plants. New Phytologist, 216, 591–604.

Furumizu, C., Alvarez, J. P., Sakakibara, K. & Bowman, J. L. 2015. Antagonistic Roles for Knox1 and Knox2 Genes in Patterning the Land Plant Body Plan Following an Ancient Gene Duplication. Plos Genetics, 11, e1004980.

Gillissen, B., Bergemann, J., Sandmann, C., Schroeer, B., Bolker, M. & Kahmann, R. 1992. A 2-Component Regulatory System for Self Non-Self Recognition in Ustilago-Maydis. Cell, 68, 647–657.

Goodrich, J., Puangsomlee, P., Martin, M., Long, D., Meyerowitz, E. M. & Coupland, G. 1997. A polycomb-group gene regulates homeotic gene expression in Arabidopsis. Nature, 386, 44–51.

Goutte, C. & Johnson, A. D. 1988. A1-Protein Alters the Dna-Binding Specificity of Alpha-2 Repressor. Cell, 52, 875–882.

Hackbusch, J., Richter, K., Muller, J., Salamini, F. & Uhrig, J. F. 2005. A central role of Arabidopsis thaliana ovate family proteins in networking and subcellular localization of 3-aa loop extension homeodomain proteins. Pnas, 102, 4908–4912.

Hay, A. & Tsiantis, M. 2010. Knox genes: versatile regulators of plant development and diversity. Development 137, 3153–3165.

Hedgethorne, K., Eustermann, S., Yang, J.-C., Ogden, T. E. H., Neuhaus, D. & Bloomfield, G. 2017. Homeodomain-like Dna binding proteins control the haploid-to-diploid transition in Dictyostelium. Science Advances, 3, e1602937.

Hellens, R. P., Edwards, E. A., Leyland, N. R., Bean, S. & Mullineaux, P. M. 2000. pGreen: a versatile and flexible binary Ti vector for Agrobacterium-mediated plant transformation. Plant Molecular Biology, 42, 819–832.

Herskowitz, I. 1989. A Regulatory Hierarchy for Cell Specialization in Yeast. Nature, 342, 749–757.

Higo, A., Niwa, M., Yamato, K. T., Yamada, L., Sawada, H., Sakamoto, T., Kurata, T., Shirakawa, M., Endo, M., Shigenobu, S., Yamaguchi, K., Ishizaki, K., Nishihama, R., Kohchi, T. & Araki, T. 2016. Transcriptional Framework of Male Gametogenesis in the Liverwort Marchantia polymorpha L. Plant and Cell Physiology, 57, 325–338.

Hisanaga, T., Okahashi, K., Yamaoka, S., Kajiwara, T., Nishihama, R., Shimamura, M., Yamato, K. T., Bowman, J. L., Kohchi, T. & Nakajima, K. 2019. A cis-acting bidirectional transcription switch controls sexual dimorphism in the liverwort. The Embo Journal, e100240.

Hofmeister, W. F. B. 1862. On the germination, development, and fructification of the higher Cryptogamia, and on the fructification of the Coniferae., London, Ray Society.

Hori, K., Maruyama, F., Fujisawa, T., Togashi, T., Yamamoto, N., Seo, M., Sato, S., Yamada, T., Mori, H., Tajima, N., Moriyama, T., Ikeuchi, M., Watanabe, M., Wada, H., Kobayashi, K., Saito, M., Masuda, T., Sasaki-Sekimoto, Y., Mashiguchi, K., Awai, K., Shimojima, M., Masuda, S., Iwai, M., Nobusawa, T., Narise, T., Kondo, S., Saito, H., Sato, R., Murakawa, M., Ihara, Y., Oshima-Yamada, Y., Ohtaka, K., Satoh, M., Sonobe, K., Ishii, M., Ohtani, R., Kanamori-Sato, M., Honoki, R., Miyazaki, D., Mochizuki, H., Umetsu, J., Higashi, K., Shibata, D., Kamiya, Y., Sato, N., Nakamura, Y., Tabata, S., Ida, S., Kurokawa, K. & Ohta, H. 2014. Klebsormidium flaccidum genome reveals primary factors for plant terrestrial adaptation. Nature Communications, 5, 3978.

Horst, N. A., Katz, A., Pereman, I., Decker, E. L., Ohad, N. & Reski, R. 2016. A single homeobox gene triggers phase transition, embryogenesis and asexual reproduction. Nature Plants, 2, 15209.

Huelsenbeck, J. P. & Ronquist, F. 2001. Mrbayes: Bayesian inference of phylogenetic trees. Bioinformatics, 17, 754–755.

Huelsenbeck, J. P., Ronquist, F., Nielsen, R. & Bollback, J. P. 2001. Evolution - Bayesian inference of phylogeny and its impact on evolutionary biology. Science, 294, 2310–2314.

Hull, C. M., Boily, M. J. & Heitman, J. 2005. Sex-specific homeodomain proteins Sxi1 alpha and Sxi2a coordinately regulate sexual development in Cryptococcus neoformans. Eukaryotic Cell, 4, 526–535.

Inoue, K., Nishihama, R., Araki, T. & Kohchi, T. 2019. Reproductive induction is a far-red high irradiance response that is mediated by phytochrome and Phytochrome Interacting Factor in Marchantia polymorpha. Plant and Cell Physiology, 60, 1136–1145.

Ishizaki, K., Chiyoda, S., Yamato, K. T. & Kohchi, T. 2008. Agrobacterium-mediated transformation of the haploid liverwort Marchantia polymorpha L., an emerging model for plant biology. Plant and Cell Physiology, 49, 1084–1091.

Ishizaki, K., Nishihama, R., Ueda, M., Inoue, K., Ishida, S., Nishimura, Y., Shikanai, T. & Kohchi, T. 2015. Development of Gateway Binary Vector Series with Four Different Selection Markers for the Liverwort Marchantia polymorpha. Plos one, 10, e0138876.

Jones, R. S. & Gelbart, W. M. 1993. The Drosophila Polycomb-Group Gene Enhancer of Zeste Contains a Region with Sequence Similarity to Trithorax. Molecular and Cellular Biology, 13, 6357–6366.

Joo, S., Wang, M. H., Liu, G., Lee, J., Barnas, A., Kim, E., Sudek, S., Worden, A. Z. & Lee, J.-H. 2018. Common ancestry of heterodimerizing Tale homeobox transcription factors across Metazoa and Archaeplastida. Bmc Biology, 16, 136.

Ju, C. L., Van De Poel, B., Cooper, E. D., Thierer, J. H., Gibbons, T. R., Delwiche, C. F. & Chang, C. R. 2015. Conservation of ethylene as a plant hormone over 450 million years of evolution. Nature Plants, 1, 14004.

Kamada, T. 2002. Molecular genetics of sexual development in the mushroom Coprinus cinereus. BioEssays, 24, 449–459.

Kariyawasam, T., Joo, S., Lee, J., Toor, D., Gao, A. F., Noh, K.-C. & Lee, J.-H. 2019. Tale homeobox heterodimer Gsm1-Gsp1 is a molecular switch that prevents unwarranted genetic recombination in Chlamydomonas. The Plant Journal, 100, 938–953.

Keeling, P. J., Burki, F., M. Wilcox, H., Allam, B., Allen, E. E., Amaral-Zettler, L. A., Armbrust, E. V., Archibald, J. M., Bharti, A. K., Bell, C. J., Beszteri, B., Bidle, K. D., T. Cameron, C., Campbell, L., Caron, D. A., Cattolico, R. A., Jackie L. Collier4, Coyne, K., Davy, S. K., Deschamps, P., Dyhrman, S. T., Edvardsen, B., Gates, R. D., Gobler, C. J., Greenwood, S. J., Guida, S. M., Jacobi, J. L., Jakobsen, K. S., James, E. R., Jenkins, B., John, U., Johnson, M. D., Juhl, A. R., Kamp, A., Katz, L. A., Kiene, R., Kudryavtsev, A., Leander, B. S., Lin, S., Lovejoy, C., Lynn, D., Marchetti, A., Mcmanus, G., Nedelcu, A. M., Menden-Deuer, S., Miceli, C., Mock, T., Montresor, M., Moran, M. A., Murray, S., Nadathur, G., Nagai, S., Ngam, P. B., Palenik, B., Pawlowski, J., Petroni, G., Piganeau, G., Posewitz, M. C., Rengefors, K., Romano, G., Rumpho, M. E., Rynearson, T., Schilling, K. B., Schroeder, D. C., Simpson, A. G. B., Slamovits, C. H., Smith, D. R., Smith, G. J., Smith, S. R., Sosik, H. M., Stief, P., Theriot, E., Twary, S. N., Umale, P. E., Vaulot, D., Wawrik, B., Wheeler, G. L., Wilson, W. H., Xu, Y., Zingone, A. & Z.Worden, A. 2014. The Marine Microbial Eukaryote Transcriptome Sequencing Project (Mmetsp): Illuminating the Functional Diversity of Eukaryotic Life in the Oceans through Transcriptome Sequencing. Plos Biology, 6, e1001889.

Khanday, I., Skinner, D., Yang, B., Mercier, R. & Sundaresan, V. 2019. A male-expressed rice embryogenic trigger redirected for asexual propagation through seeds. Nature, 565, 91–95.

Kienitz-Gerloff, F. 1874. Vergleichende Untersuchungen über die Entwickelungsgeschichte des Lebermoos - Sporogoniums. Botanische Zeitung, 32, 161–172; 193-204; 209-221; 225-238.

Kimura, S., Koenig, D., Kang, J., Yoong, F. Y. & Sinha, N. 2008. Natural variation in leaf morphology results from mutation of a novel Knox gene. Current Biology, 18, 672–677.

Knapp, E. 1935. Untersuchungen über die Wirkung von Röntgenstrahlen an dem Lebermoos Sphaerocarpus, mit Hilfe der tetraden-Analyse. I. Zeitschrift für Induktive Abstammungs- und Vererbungslehre, 70, 309–349.

Kues, U., Richardson, W. V. J., Tymon, A. M., Mutasa, E. S., Gottgens, B., Gaubatz, S., Gregoriades, A. & Casselton, L. A. 1992. The Combination of Dissimilar Alleles of the a-Alpha and a-Beta Gene Complexes, Whose Proteins Contain Homeo Domain Motifs, Determines Sexual Development in the Mushroom Coprinus-Cinereus. Genes & Development, 6, 568–577.

Lampugnani, E. R., Ho, Y. Y., Moller, I. E., Koh, P.-L., Golz, J. F., Bacic, A. & Newbigin, E. 2016. A glycosyltransferase from Nicotiana alata pollen mediates synthesis of a linear (1, 5)-α-L-arabinan when expressed in Arabidopsis. Plant physiology, 170, 1962–1974.

Lee, J.-H., Lin, H., Joo, S. & Goodenough, U. 2008. Early Sexual Origins of Homeoprotein Heterodimerization and Evolution of the Plant Knox/Bell Family. Cell, 133, 829–840.

Lewis, E. B. 1978. A gene complex controlling segmentation in Drosophila. Nature, 276, 565–570.

Li, H. & Durbin, R. 2009. Fast and accurate short read alignment with Burrows-Wheeler transform. Bioinformatics, 25, 1754–1760.

Lorbeer, G. 1936. Die Umwandlung eines haploiden, genotypisch weiblichen Gametophyten von Sphaerocarpus Donnellii in einen männlichen mit Hilfe von Röntgenstrahlen. Planta, 25, 70–83.

Magnani, E. & Hake, S. 2008. Knox lost the Ox: The Arabidopsis Knatm gene defines a novel class of Knox transcriptional regulators missing the homeodomain. Plant Cell, 20, 875–887.

Marchant, J. 1713. Nouvelle Découverte des fleurs et des graines D’une Plante rangée par les Botanistes fous le genre du Lichen. Histoire de l’Académie royale des sciences, avec les mémoires de mathématique et de physique, pour la meme cette Académie., Année Mdccxiii, 229-234.

Merabet, S. & Mann, R. S. 2016. To Be Specific or Not: The Critical Relationship Between Hox And Tale Proteins. Trends in Genetics, 32, 334–347.

Meyberg, R., Perroud, P.-F., Haas, F. B., Schneider, L., Heimerl, T., Renzaglia, K. & Rensing, S. A. 2020. Characterization of evolutionarily conserved key players affecting eukaryotic flagellar motility and fertility using a moss model. New Phytologist, doi.org/10.1111/nph.16486.

Mikami, K., Li, C., Irie, R. & Hama, Y. 2019. A unique life cycle transition in the red seaweed Pyropia yezoensis depends on apospory. Communications Biology, 2, 299.

Mirbel, C.-F. 1835. Researches anatomiques et physiologiques sur le Marchantia polymorpha. (Anatomical and physiological researches on Marchantia polymorpha.). Mém. Acad. Roy. Sc. Inst. France., 13, 337–436

Montgomery, S. A., Tanizawa, Y., Galik, B., Wang, N., Ito, T., Mochizuki, T., Akimcheva, S., Bowman, J. L., Cognat, V., Maréchal-Drouard, L., Ekker, H., Hong, S.-F., Kohchi, T., Lin, S.-S., Liu, L.-Y. D., Nakamura, Y., Valeeva, L. R., Shakirov, E. V., Shippen, D. E., Wei, W.-L., Yagura, M., Yamaoka, S., Yamato, K. T., Liu, C. & Berger, F. 2020. Chromatin Organization in Early Land Plants Reveals an Ancestral Association between H3K27me3, Transposons, and Constitutive Heterochromatin. Current Biology, 30, 573–588.e7.

Mosquna, A., Katz, A., Decker, E. L., Rensing, S. A., Reski, R. & Ohad, N. 2009. Regulation of stem cell maintenance by the Polycomb protein Fie has been conserved during land plant evolution. Development, 136, 2433–2444.

Muller, J., Hart, C. M., Francis, N. J., Vargas, M. L., Sengupta, A., Wild, B., Miller, E. L., O’connor, M. B., Kingston, R. E. & Simon, J. A. 2002. Histone methyltransferase activity of a Drosophila polycomb group repressor complex. Cell, 111, 197–208.

Nishimura, Y., Shikanai, T., Nakamura, S., Kawai-Yamada, M. & Uchimiya, H. 2012. Gsp1 Triggers the Sexual Developmental Program Including Inheritance of Chloroplast Dna and Mitochondrial Dna in Chlamydomonas reinhardtii. Plant Cell, 24, 2401–2414.

Okano, Y., Aonoa, N., Hiwatashia, Y., Murata, T., Nishiyama, T., Ishikawa, T., Kubo, M. & Hasebe, M. 2009. A polycomb repressive complex 2 gene regulates apogamy and gives evolutionary insights into early land plant evolution. Pnas, 106, 16321–26.

Ortiz-Ramírez, C., Michard, E., Simon, A. A., Damineli, D. S. C., Hernández-Coronado, M., Becker, J. D. & Feijo, J. A. 2017. Glutamate Receptor-Like channels are essential for chemotaxis and reproduction in mosses. Nature, 549, 91–95.

Pagnussat, G. C., Yu, H. J. & Sundaresana, V. 2007. Cell-fate switch of synergid to egg cell in Arabidopsis eostre mutant embryo sacs arises from misexpression of the Bel1-like homeodomain gene Blh1. Plant Cell, 19, 3578–3592.

Pearson, J. C., Lemons, D. & Mcginnis, W. 2005. Modulating Hox gene functions during animal body patterning. Nature Reviews Genetics, 6, 893–904.

Pereman, I., Mosquna, A., Katz, A., Wiedemann, G., Lang, D., Decker, E. L., Tamada, Y., Ishikawa, T., Nishiyama, T., Hasebe, M., Reski, R. & Ohad, N. 2016. The Polycomb group protein Clf emerges as a specific tri-methylase of H3K27 regulating gene expression and development in Physcomitrella patens. Biochimica Et Biophysica Acta-Gene Regulatory Mechanisms, 1859, 860–870.

Renzaglia, K. S., Villarreal, J. C. & Garbary, D. J. 2018. Morphology supports the setaphyte hypothesis-mosses plus liverworts form a natural group. Bryophyte Diversity & Evolution, 40, 11–17.

Retamales, H. A. & Scharaschkin, T. 2014. A staining protocol for identifying secondary compounds in Myrtaceae. Applications in Plant Sciences, 2, 1400063.

Sakakibara, K. 2016. Technological Innovations Give Rise to a New Era of Plant Evolutionary Developmental Biology. *In*: Rensing, S. A. (ed.) Advances in Botanical Research: Genomes and Evolution of Charophytes, Bryophytes, Lycophytes and Ferns. Elsevier.

Sakakibara, K., Ando, S., Yip, H. K., Tamada, Y., Hiwatashi, Y., Murata, T., Deguchi, H., Hasebe, M. & Bowman, J. L. 2013. Knox2 genes regulate the haploid-to-diploid morphological transition in land plants. Science, 339, 1067–1070.

Sakakibara, K., Nishiyama, T., Deguchi, H. & Hasebe, M. 2008. Class 1 Knox genes are not involved in shoot development in the moss *Physcomitrella patens* but do function in sporophyte development. Evol. Dev., 10, 555–566.

Sano, R., Juarez, C. M., Hass, B., Sakakibara, K., Ito, M., Banks, J. A. & Hasebe, M. 2005. Knox homeobox genes potentially have similar function in both diploid unicellular and multicellular meristems, but not in haploid meristems. Evolution & Development, 7, 69–78.

Saurin, A. J., Shao, Z. H., Erdjument-Bromage, H., Tempst, P. & Kingston, R. E. 2001. A Drosophila Polycomb group complex includes Zeste and dtafii proteins. Nature, 412, 655–660.

Schulz, K. N. & Harrison, M. M. 2019. Mechanisms regulating zygotic genome activation. Nature Reviews Genetics, 20, 221–234.

Singer, S. D. & Ashton, N. W. 2007. Revelation of ancestral roles of Knox genes by a functional analysis of Physcomitrella homologues. Plant Cell Reports, 26, 2039–2054.

Smith, H. M. S., Boschke, I. & Hake, S. 2002. Selective interaction of plant homeodomain proteins mediates high Dna-binding affinity. Pnas, 99, 9579–9584.

Spit, A., Hyland, R. H., Mellor, E. J. C. & Casselton, L. A. 1998. A role for heterodimerization in nuclear localization of a homeodomain protein. Pnas, 95, 6228–6233.

Sugano, S. S., Nishihama, R., Shirakawa, M., Takagi, J., Matsuda, Y., Ishida, S., Shimada, T., Hara-Nishimura, I., Osakabe, K. & Kohchi, T. 2018. Efficient Crispr/Cas9-based genome editing and its application to conditional genetic analysis in Marchantia polymorpha. PloS One, 13, e0205117.

Sugano, S. S., Shirakawa, M., Takagi, J., Matsuda, Y., Shimada, T., Hara-Nishimura, I. & Kohchi, T. 2014. Crispr/Cas9-Mediated Targeted Mutagenesis in the Liverwort Marchantia polymorpha L. Plant and Cell Physiology, 55, 475–481.

Swanson, W. J. & Vacquier, V. D. 2002. The rapid evolution of reproductive proteins. Nature Reviews Genetics, 3, 137–144.

Thuret, G. 1851. Recherches sur les Zoospores des Algues et les Anthéridies des Cryptogames. Annales de Sciences Naturelles Botanique, 3.16, 5–39.

Unger, F. 1837. Weitere Beobachtungen über die Samenthiere der Pflanzen. Nova Acta physico medica A C L, 18, 786–796.

Urban, M., Kahmann, R. & Bolker, M. 1996. Identification of the pheromone response element in Ustilago maydis. Molecular & General Genetics, 251, 31–37.

Vanneste, K., Sterck, L., Myburg, A. A., Van De Peer, Y. & Mizrachi, E. 2015. Horsetails Are Ancient Polyploids: Evidence from Equisetum giganteum. Plant Cell, 27, 1567–1578.

Wickett, N. J., Mirarab, S., Nguyen, N., Warnow, T., Carpenter, E., Matasci, N., Ayyampalayam, S., Barker, M. S., Burleigh, J. G., Gitzendanner, M. A., Ruhfel, B. R., Wafula, E., Der, J. P., Graham, S. W., Mathews, S., Melkonian, M., Soltis, D. E., Soltis, P. S., Miles, N. W., Rothfels, C. J., Pokorny, L., Shaw, A. J., Degironimo, L., Stevenson, D. W., Surek, B., Villarreal, J. C., Roure, B., Philippe, H., Depamphilis, C. W., Chen, T., Deyholos, M. K., Baucom, R. S., Kutchan, T. M., Augustin, M. M., Wang, J., Zhang, Y., Tian, Z. J., Yan, Z. X., Wu, X. L., Sun, X., Wong, G. K. S. & Leebens-Mack, J. 2014. Phylotranscriptomic analysis of the origin and early diversification of land plants. Pnas, 111, E4859–E4868.

Zachgo, S. 2002. In situ hybridization. *In*: Gilartin, P. M. & Blower, C. (eds.) Molecular Plant Biology. Oxford, Uk: Oxford University Press.

Zhang, Y., Liu, T., Meyer, C. A., Eeckhoute, J., Johnson, D. S., Bernstein, B. E., Nussbaum, C., Myers, R. M., Brown, M., Li, W. & Liu, X. S. 2008. Model-based Analysis of Chip-Seq (Macs). Genome Biology, 9, R137.

Zhao, H., Lu, M., Singh, R. & Snell, W. J. 2001. Ectopic expression of a Chlamydomonas mt+-specific homeodomain protein in mt− gametes initiates zygote development without gamete fusion. Genes & Development, 15, 2767–2777.

